# Adaptive Machine Learning Framework enables Unprecedented Yield and Purity of Adeno-Associated Viral Vectors for Gene Therapy

**DOI:** 10.1101/2025.05.23.655859

**Authors:** Kelvin P. Idanwekhai, Shriarjun Shastry, Arianna Minzoni, Morgan R. Hurst, Eduardo Barbieri, Eugene N. Muratov, Michael A. Daniele, Stefano Menegatti, Alexander Tropsha

**Author notes:** Authors contributed equally (Kelvin P. Idanwekhai, Shriarjun Shastry, Arianna Minzoni).

## Abstract

Adeno-associated viral (AAV) vectors for gene therapy are becoming integral to modern medicine, providing therapeutic options for diseases once deemed incurable. Currently, optimizing viral vector purification is a critical bottleneck in the gene therapy industry, impacting product efficacy and safety as well as accessibility and cost to patients. Traditional optimization methods are resource-intensive and often fail to adjust the purification process parameters to maximize the resulting product yield and quality. To address this challenge, we developed a machine learning framework that leverages Bayesian optimization to systematically refine affinity chromatography parameters (sample load, flow rate, and the formulation of chromatographic media) to improve AAV purification. The efficiency of this closed-loop workflow in iteratively optimizing the vector’s yield, purity, and transduction efficiency was demonstrated by purifying clinically-relevant serotypes AAV2, AAV5, and AAV9 from HEK293 cell lysates using the affinity adsorbent AAVidity. We show that three cycles of Bayesian optimization elevated yields from a baseline of 70% to 99%, while reducing host-cell impurities by 230-to-400-fold across all serotypes. The optimized parameters consistently produced vectors with high purity and preserved high transduction activity, essential for therapeutic efficacy and safety, demonstrating serotype versatility – a key challenge in AAV manufacturing. By streamlining parameter optimization and enhancing productivity, our adaptive machine learning framework accelerates process development and reduces costs, advancing the accessibility and clinical translation of AAV-based gene therapies.

## 1. Introduction

Adeno-associated viruses (AAVs) have become the leading delivery platform technology for in vivo gene therapies owing to their high transduction efficiency, broad tissue tropism, and low immunogenicity^1–5^. The design of novel AAVs and genetic payloads with improved therapeutic efficacy and safety is progressing rapidly, fueled by the recent advancements in synthetic biology and artificial intelligence-guided design of viral capsids^2,5–9^. On the other hand, the technologies for industrial biomanufacturing of AAVs, particularly in the enrichment and purification of functional virions, have not kept pace. The current AAV purification process comprises two chromatographic steps, namely product capture by affinity chromatography, which separates AAVs from host cell proteins and DNA, followed by product polishing by ion-exchange chromatography, which separates gene-loaded capsids from empty or partially loaded particles and removes residual impurities^10–14^. The unique design of each gene therapy product – ground in the combination of AAV serotype and genetic payload – produces variability in virion titer, capsid content ratios, and impurity profiles, necessitating rigorous optimization of process parameters at both capture and polishing stages. Current optimization strategies, based on simple statistical methods, often fail to capture the complex, multivariate relationships linking feed composition, AAV-ligand interactions, and mobile phase properties to the process outcomes, resulting in suboptimal yields, purity, and transduction activity^13,15–17^. These inefficiencies contribute to high production costs, limited availability, and, in some cases, adverse clinical outcomes that have impacted public confidence in gene therapies^3,18,19^. The space of input process parameters for chromatographic optimization is vast, making exhaustive exploration via conventional approaches-such as mechanistic modeling, Design of Experiments (DoE), and hybrid models, impractical. Mechanistic models provide detailed insights into fluid dynamics and mass transport, but are challenging to calibrate due to the need for extensive parameterization^20–24^. DoE methods, such as Response Surface Methodology (RSM), are easier to implement but often lack predictive power outside the tested design space^25–27^. Hybrid models combine mechanistic and empirical elements to reduce experimental burden while maintaining predictive capability, yet they struggle with the high-dimensional data typical of AAV purification^28–30^.

Data-driven machine learning approaches represent a transformative toolkit for accelerating the optimization of viral vector purification processes. Bayesian optimization (BO) has emerged as a particularly effective strategy for globally maximizing complex, non-linear systems with high experimental costs, such as affinity chromatography, where fluid dynamics, mass transfer, and ligand-capsid interactions intersect^31–34^. This methodology employs probabilistic machine learning (ML) models to strategically select experimental parameters, continuously refining predictions through iterative, model-guided data acquisition^35–38^. By balancing the exploration of the design space with the exploitation of accumulated knowledge, BO efficiently navigates noisy, non-convex optimization landscapes even with sparse datasets^35,38,39^. Recent studies validated ML-guided closed-loop experimental systems and autonomous laboratory platforms for process development^40–45^, demonstrating their applicability to biopharmaceutical manufacturing.

In this study, we implemented an adaptive BO framework that integrates a Gaussian Process and a *qExpectedImprovement(qEI)* acquisition function to establish a closed-loop optimization system for AAV affinity chromatography (**Figure 1**). The workflow iteratively updates purification parameters based on prior experimental outcomes until achieving target yield and purity thresholds, with each optimization cycle constrained to ten experimental trials. We demonstrated the effectiveness of this bioprocess development platform by optimizing the purification of three clinically relevant serotypes –AAV2, AAV5, and AAV9 – from HEK293 cell lysates using the affinity adsorbent AAVidity^46–48^. To evaluate the transferability of our approach, we used the data acquired with AAV2 to seed the optimization of the AAV9 purification process.

**Figure 1.**
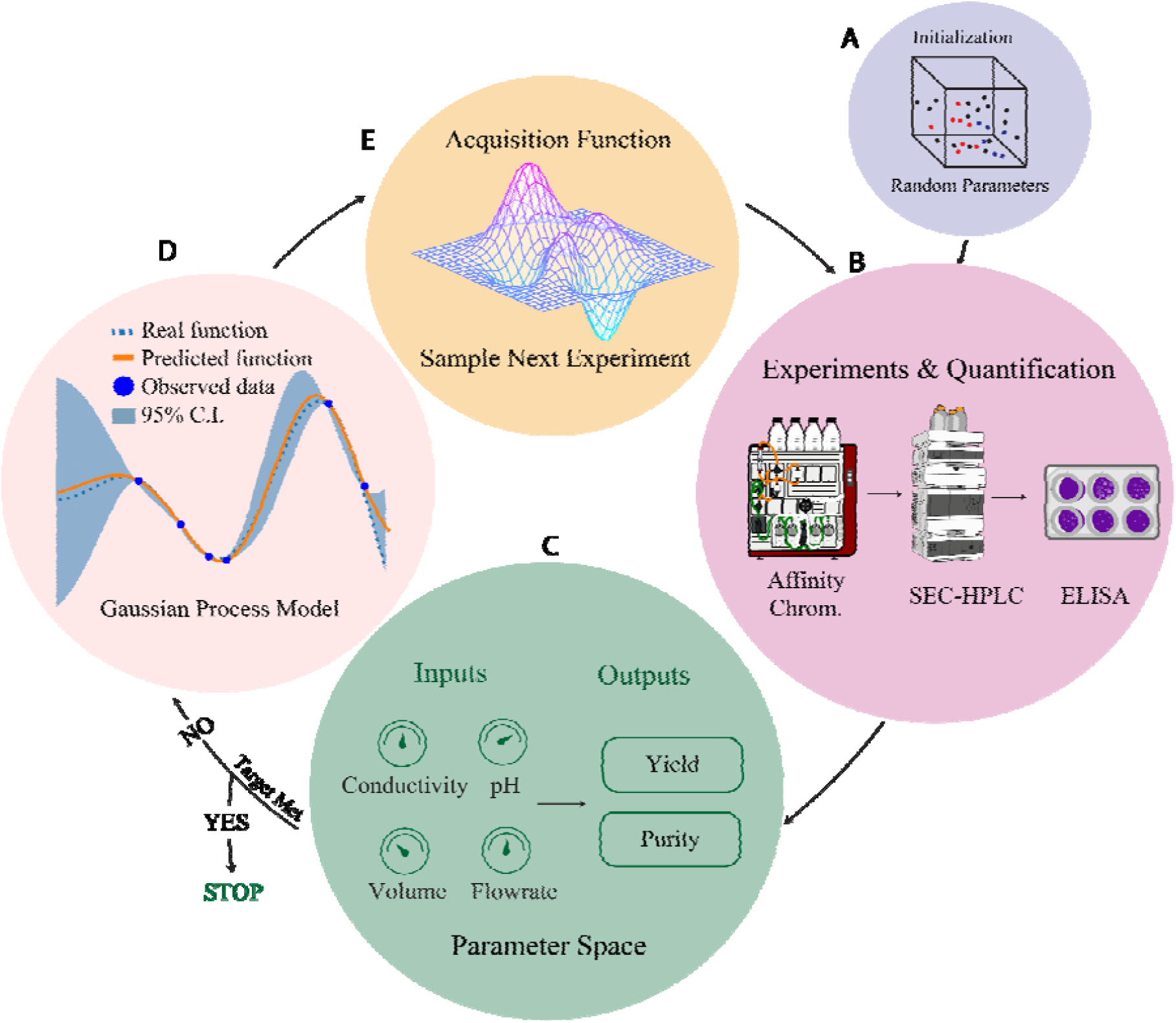
Closed-loop optimization of AAV purification by affinity chromatography: **(A)** selection of an initial random set of process input parameters; **(B)** experimental data collection; **(C)** purification performance results recorded in step B are used for model development; **(D)** build and update a Gaussian process surrogate model using input and output process parameters; **(E)** implement an acquisition function to select the values of input parameter for the subsequent chromatography test set. The proposed optimization cycle iterates steps (B), (C), (D), and (E) until the optimization objectives (i.e., AAV yield and purity) are reached.

We find that the BO-driven workflow succeeded in identifying optimal elution conditions within limited iterations, achieving over 95% yield and 99% purity across all serotypes. Unlike conventional methods that prioritize yield at the expense of quality metrics, our platform concurrently optimized capsid recovery and host cell protein removal while safeguarding the transduction activity of the purified vectors. We posit that this novel optimization methodology can be applied to a broad host of AAV to ensure therapeutic efficacy while maintaining compliance with regulatory requirements for gene therapy products.

## 2. Results and Discussion

### 2.1. Closed-loop experimentation design and method development

The multidimensional and complex nature of interconnected process parameters in AAV affinity chromatography limits the effectiveness of conventional optimization methodologies for achieving substantial improvements in product yield and quality. To address this challenge, we implemented an adaptive optimization framework employing BO to iteratively refine process parameters through experimental validation. This closed-loop system, outlined in **Figure 1**, updates a probabilistic surrogate model with purification outcomes, enabling data-driven adjustments to subsequent experimental conditions.

The initial model validation focused on the purification of the AAV “legacy” serotype 2 (AAV2), with viral capsid yield as the primary optimization target. The optimization campaign was iterated by expanding from single to a dual objective (yield and purity), although the single-objective optimization alone afforded excellent outcomes. To demonstrate serotype-independent applicability, we implemented the model on AAV5 and AAV9, two clinically prominent serotypes with distinct bioprocessing challenges, thus demonstrating cross-serotype transferability of process knowledge and rapid process development.

Model development was initialized by leveraging a historical data built by purifying seven AAV serotypes (AAV2, 3, 5, 6, 8, 9, and rh.10) using the affinity resin AAVidity^46–48^, and encompassing critical process parameters such as load ratio (capsids/mL resin), flow rates, and the formulation of the chromatographic buffers used for resin equilibration and washing, and AAV elution. Unlike conventional affinity resins, AAVidity is functionalized with peptide ligands whose affinity for active, gene-loaded capsids can be adjusted by modulating a number of process parameters to meet specific process needs. The parameters extracted from the historical purification records were mapped to the resulting values of AAV titer and purity in the eluted products (**Figure S1**). The dataset was curated and randomly partitioned into a larger subset of 160 purification tests, which was used to train data regression models, including Gaussian Process (GP), random forest (RF), gradient boosting (XG Boost), and support vector machines (SVM), and a smaller hold-out set of 40 tests, which was utilized for model validation. To evaluate the accuracy of the model for predicting capsid titer based on input process parameters, we measured the coefficient of determination (R^2^) and the root mean squared error (RMSE) metrics for each regression model (**Figure S2** and **Table 1**). The GP regression model showed the highest R^2^ and the lowest RMSE of 0.92 and 1.13, respectively. Gaussian Process is desirable for bioprocess applications, which process high-value products, because it can directly provide uncertainty estimates for each prediction unlike other approaches where models must be combined in an ensemble to obtain a confidence score^49^. The GP model structure, defined by its mean *m(x)* and covariance function *k(x, x’)*, also known as kernel, follows the form ln[TCT(*x*)] ∼ GP(*m(x), k(x,x’)*), wherein TCT(*x*) is the total capsid titer obtained in response to input parameters *x*. The kernel used for the prediction model ln[TCT(*x*)] is described in *Supplementary Information* (*Section SX*).

**Table 1.** Evaluation of the data regression models trained to predict the total AAV capsid titer.

| Model | MSE | MAE | RMSE | $R^2$ |
| --- | --- | --- | --- | --- |
| XG Boost | 1.97 | 0.91 | 1.4 | 0.88 |
| Random Forest | 1.66 | 0.88 | 1.29 | 0.9 |
| SVM | 1.95 | 1.07 | 1.4 | 0.88 |
| Gaussian Process | 1.29 | 0.83 | 1.13 | 0.92 |

### 2.2. Space of design of chromatographic processes and simulation of an optimization campaign

Having validated a GP model that accurately correlates input process parameters with resultant product yield, we proceeded to implement Bayesian optimization to guide the purification of AAVs by affinity chromatography. Among the input parameters, certain variables were held constant, while others were systematically varied, either as discrete or continuous variables, to constrain the acquisition function’s sampling to experimentally feasible regions. The parameters maintained at fixed values included a solvent conductivity of 2.5 mS/cm for resin equilibration prior to feedstock loading and a standard linear velocity of 306 cm/hr during both post-loading resin wash and capsid elution. These values were identified as optimal for maximizing the dynamic binding capacity and product capture^50–53^. The variable parameters included the target AAV serotype (AAV2, AAV5, and AAV9), load volume (5 to 30 mL), linear velocity during wash and elution steps (130 to 600 cm/h), buffer pH (5.0 to 9.0), and buffer conductivity (1 to 15 mS/cm for wash and 10 to 100 mS/cm for elution). The incremental steps for each parameter were selected to enable a granular exploration of the selected ranges. The resulting process design space comprised over 930,000 unique experimental conditions, rendering exhaustive empirical investigation impractical and underscoring the necessity for intelligent, targeted exploration. To further rationalize the search space, specific pH and conductivity ranges for elution buffers were assigned for each target AAV serotype.

To estimate the duration of the optimization campaign – namely, the number of experimental iterations required to identify process parameters that maximize product yield – we conducted simulations evaluating several heuristic acquisition functions: qExpectedImprovement (*qEI*), qLogExpectedImprovement (*qLogEI*), qProbabilityofImprovement (*qPI*), and qUpperConfidenceBounds (*qUCB*). The adoption of an appropriate acquisition function, which is critical for effective optimization, was informed by considerations of process variability, measurements noise, and multi-objective optimization criteria^54–58^. The performance of each acquisition function was assessed to ensure their capacity to drive meaningful improvements in capsid yield maximization. Initial random sampling of the parameter space (**Tables S1** and **S2**) established a baseline, after which the acquisition functions were evaluated using quasi-Monte Carlo (qMC) sampling to iteratively identify parameter arrays (*x’*) that maximize the objective function ln[TCT(*x*)] (**Equation 5**) ^59,60^. To emulate a realistic experimental context, we utilized a model trained on experimental data as a source of ground truth for the simulation.

All qMC-based acquisition strategies outperformed random sampling, achieving convergence at levels corresponding to an approximately 150-fold (e^5^-fold) improvement in total AAV capsid titer (**Figure 2**). Notably, the *qEI* function required only 30 experimental iterations to identify process parameters that maximized AAV titer in the eluate (primary optimization objective), and therefore, it was adopted as the acquisition function for our Bayesian optimization campaign.

**Figure 2.**
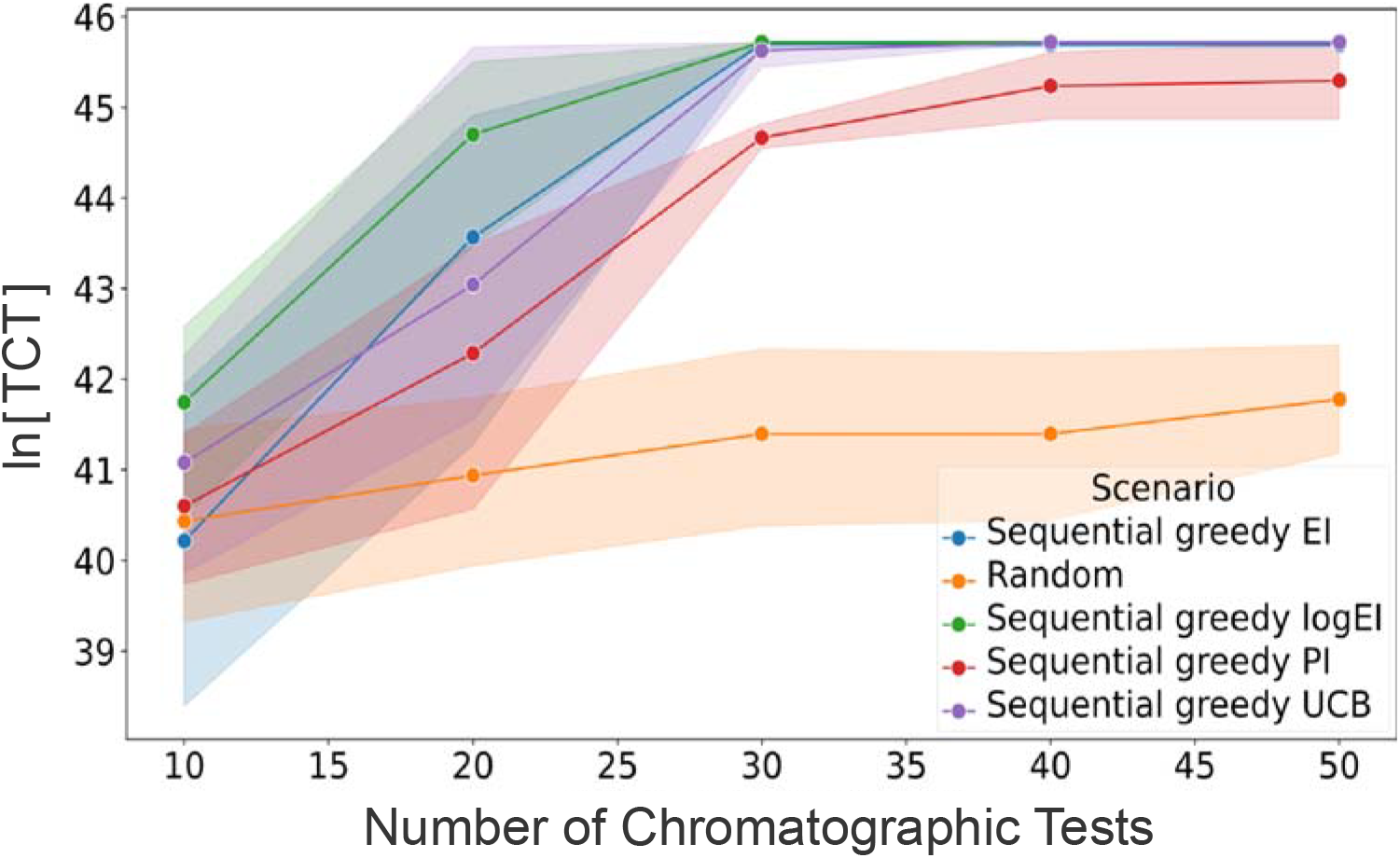
Computational simulation of the optimization campaign. Sequential acquisition functions qEI, qlogEI, qPI, and qUCB were used to simulate an experimental optimization of total capsids (TC). Simulations showed that the acquisition functions could find optimal parameters in 30 experiments, significantly surpassing a random sampling policy.

### 2.3. Bayesian optimization of AAV2, AAV5, and AAV9 purification by affinity chromatography

We employed the adaptive experimental workflow depicted in **Figure 1** to develop affinity chromatography protocols for purifying AAV2, AAV5, and AAV9 from HEK293 cell lysates. Each iteration of the Bayesian optimization cycle comprised 10 unique affinity chromatography experiments, each followed by the analysis of the eluted fractions and quantification of purification outcomes, specifically AAV yield and purity. To expedite the transition between experimental sets, we established a high-throughput assay based on analytical size exclusion chromatography (SEC-HPLC), enabling rapid assessment of AAV titer and host cell impurity clearance in eluted samples (**Figures S3-S5**). The resulting datasets of input process parameters and corresponding purification metrics were analyzed by the GP model and the *qEI* acquisition function to inform the design of subsequent experimental conditions. The initial set of ten affinity chromatography experiments for AAV2 and AAV5 was generated by random sampling of the process parameter space defined in **Tables S1** and **S2**. In contrast, the starting conditions for the AAV9 campaign were selected using the *qEI* acquisition function by leveraging data from the full history of AAV2 purification. The process input parameters utilized across all AAV purification campaigns are summarized in **Tables S3 - S5**. As shown in **Figure 3**, the progression of AAV yield values across successive Bayesian optimization cycles demonstrates a consistent enhancement in purification performance. These results indicate that, compared to conventional development approaches, our methodology achieves optimal affinity chromatography performance in thirty experiments or fewer, consistent with our simulation predictions.

**Figure 3.**
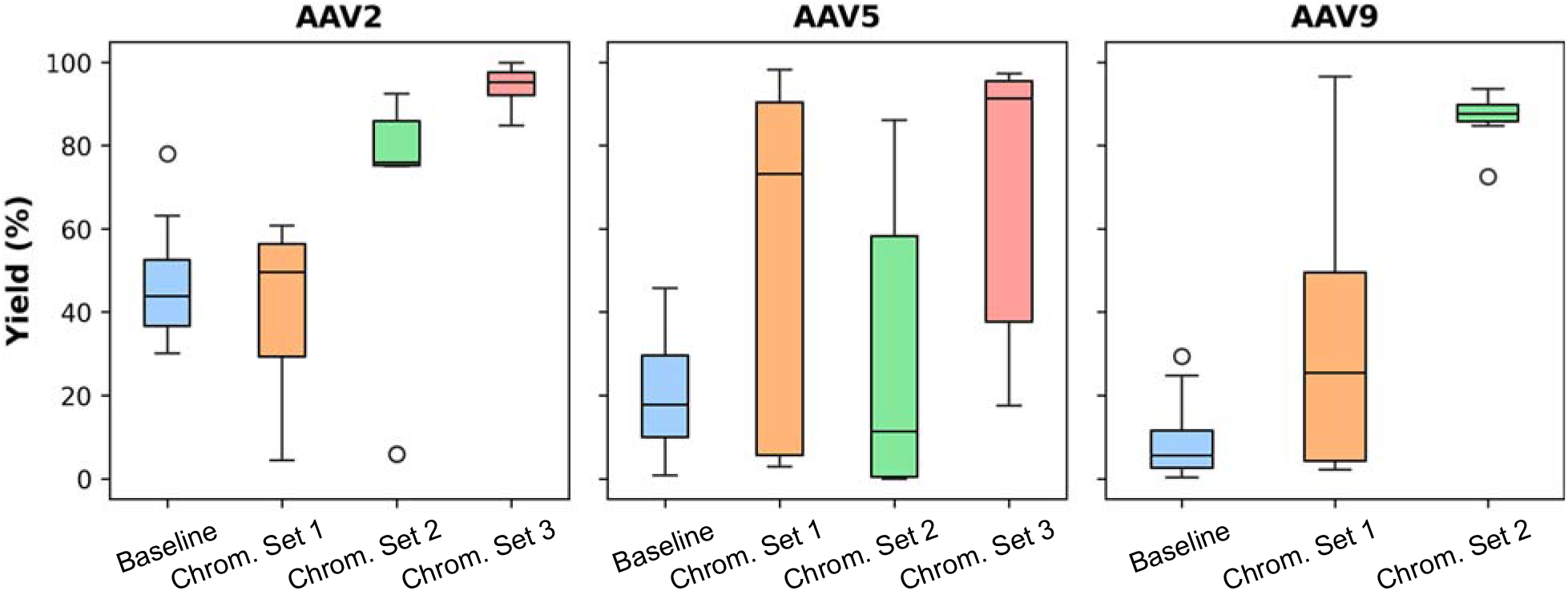
Model-guided optimization of AAV affinity chromatography demonstrates the maximization of AAV yield compared to baseline values obtained without model guidance. Comparison of product yields obtained by purifying AAV2, AAV5, and AAV9 from HEK293 cell lysates using the affinity resin AAVidity operated by machine learning-guided input process parameters.

#### 2.3.1. Purification of AAV2

Adeno-associated virus serotype 2 (AAV2), often referred to as the “legacy serotype”, is one of the most extensively studied and utilized viral vectors in gene therapy. Its broad tissue tropism facilitates efficient gene delivery to liver, muscle tissue, lungs, central nervous system (CNS), kidney, and retina^2,61–63^. Several FDA-approved gene therapies utilize AAV2 vectors, such as Luxturna for retinal dystrophy and Kebilidi for aromatic L-amino acid decarboxylase (AADC) deficiency, demonstrating its clinical significance^64–67^. Moreover, the number of active AAV2-based clinical trials reflect its continued role in advancing gene therapy. The affinity purification of AAV2 is also among the most widely documented in the chromatographic literature on viral vectors. The values of viral capsid yield achieved with commercial affinity resins such as Poros CaptureSelect AAVX and AAVidity vary between 45% and 80%^46,48,68–70^. These yields are significantly lower than those achieved with established protein therapeutics, such as monoclonal antibodies purified by Protein A affinity resins, which can reach up to 95%. Increasing AAV2 yield beyond current benchmarks could provide a significant contribution towards reducing the cost of approved gene therapies and supporting ongoing clinical evaluations.

To seed process optimization, the initial set of ten chromatographic experiments was designed by random sampling across the operational parameter space (**Table S1**), yielding an average recovery of 61%, which aligns with values reported in the literature (**Figure S3** and **Table S6**). The outcomes of this initial set were processed by the predictive model, which, guided by the *qEI* acquisition function, informed the design of a second set of ten experiments. The average yield in the second test set increased to 81% (**Figure 4A**), although two runs produced low yields (∼6%). The latter values can be attributed to the explore-exploit nature of the *qEI* function, which prioritizes the space of optimized parameters, but also attempts to explore new input parameters, where, despite the promise, the yield may fall. Leveraging the combined data from the first two sets, the *qEI* function proposed a third set of experiments, all of which achieved yields exceeding 85%. Concurrently, AAV2 purity, measured by the high-throughput SEC-HPLC assay, improved from 86-99% in the first set, to 90-99% in the second, reaching 94-99% in the third (Figure 4A). The concurrent rise in yield and purity – despite yield being the primary optimization target – suggests a synergistic relationship between these metrics.

**Figure 4.**
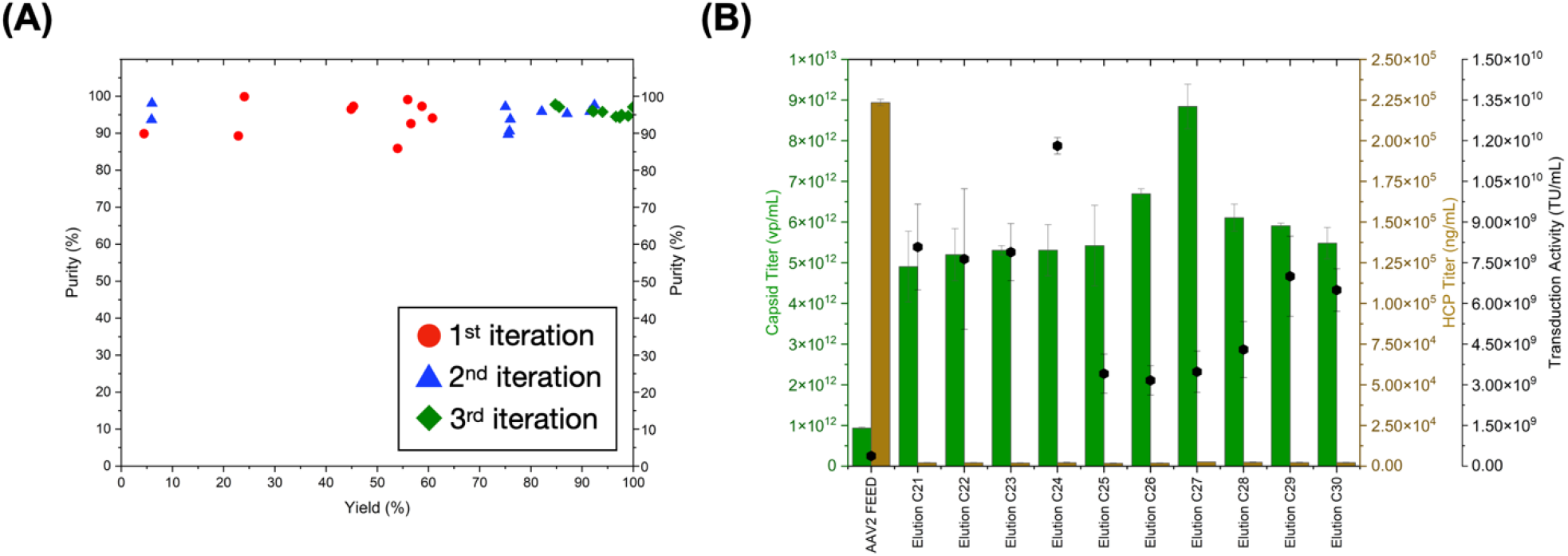
**(A)** Values of capsid yield and purity measured by SEC-HPLC analysis of the elution fractions obtained by purifying AAV2 from HEK293 cell lysates using AAVidity. Process input parameters were selected across three iterations of the closed-loop Bayesian optimization process. **(B)** Values of capsid titer, HEK293 HCP titer, and HT1080 cell transduction activity in the elution fractions obtained by purifying AAV2 using AAVidity resin operated with process parameters selected in the third iteration of Bayesian optimization.

The SEC-HPLC analysis, while providing a rapid measurement of AAV titer and purity, is not viewed as standard in the gene therapy industry. Capsid-specific ELISA kits are preferred for titer quantification, and HEK293 ELISA kits are utilized for host cell protein (HCP) measurement. Furthermore, capsid titer alone does not describe the activity of viral vectors, since not all capsids are competent for cell transduction. Therefore, selected fractions from the third chromatographic test set were further analyzed using ELISA for AAV2 titer and HCP removal, and a cell-based transduction assay in HT1080 cells to assess vector potency (**Figure 4B**). The ELISA-derived capsid titers, ranging from 4·10^12^ to 9·10^12^ vp/mL, were consistent with SEC-HPLC results and corresponded to a 5-to-9-fold enrichment over the initial feedstock. The affinity purification also achieved up to 200-fold reduction in HEK293 HCP content, from approximately 223.4 μg/mL to 1.9-2.5 μg/mL, corresponding to ∼750 ng per Luxturna-equivalent dose (∼10^11^ vg), a level admissible in clinical application. Equally as remarkable was the high yield cell-transducing AAV2 units, whose enrichment reached upwards of 1.5-fold, owing to the gentle elution conditions enabled by the AAVidity resin and selected by the BO process.

#### 2.3.2. Purification of AAV9

Among the emerging viral vectors in gene therapy, AAV9 occupies preeminent place owing to its ability to deliver genetic payloads to the central nervous system (*i.e*., brain and spinal cord)^71–74^. Numerous clinical trials have demonstrated the superior transduction efficiency of AAV9 in neuronal tissues, thus promoting its adoption in gene therapies for disorders such as spinal muscular atrophy (SMA) and muscular dystrophies, and culminating in the 2019 approval of Zolgensma as a cure for SMA^75–79^. The unique structural features of AAV9 mandate the use of serotype-specific affinity resins for effective purification since the serotype-agnostic adsorbents fail to bind AAV9 efficiently. For instance, the HiTrap AVB sepharose resin binds serotypes 2, 5, and 8, but does not capture AAV9; similarly, the Poros CaptureSelect AAVX resin does not capture AAV9 consistently from different feedstocks. To overcome these limitations, affinity resins such as CaptureSelect AAV9 and AVIPure AAV9 have been engineered specifically for AAV9 purification. In contrast, the AAVidity resin used in this study binds AAV9 under conditions and with affinity comparable to those of AAV2 binding. Accordingly, the purification outcomes from the single-objective optimization of AAV2 were used to initialize the optimization campaign of AAV9 purification.

Within the initial set of ten purification experiments designed using the *qEI* acquisition function for AAV9, five resulted in low capsid yields (2.3% – 6.5%), four achieved moderate yields (44.3% – 64.5%), whereas one achieved a notably high yield of 96.7% (**Figure S4** and **Table S7**). The suboptimal yields were attributed to AAV9’s sensitivity to elevated buffer conductivity during the post-load resin wash step. The chromatographic tests that employed wash buffers with conductivities above 9 mS/cm consistently delivered poor yields, whereas the highest-yielding condition employed a mild wash buffer (∼1 mS/cm). The susceptibility of AAV9 to high conductivity solvents is likely due to the unique charge distribution on the virion surface and the disruption of electrostatic interactions between capsid proteins, ultimately compromising capsid integrity. The *qEI* acquisition function rapidly adapted by selecting low-conductivity buffers for the subsequent set of chromatographic tests; at the same time, the model pursued the exploration of conductivities up to 4 mS/cm. The outcomes of the second chromatographic test set rewarded the choices of the acquisition function, with yields ranging from 85% to 94% (except for a 73% outlier), and values of purity averaging 99%. As anticipated, the highest-conductivity wash buffer (4 mS/cm) produced the lowest yield (72.6%), while wash buffers at 2 and 3 mS/cm yielded an average of 85.5% and 84.7%, respectively, supporting the exploration of a broader conductivity range (**Figure 5A**).

**Figure 5.**
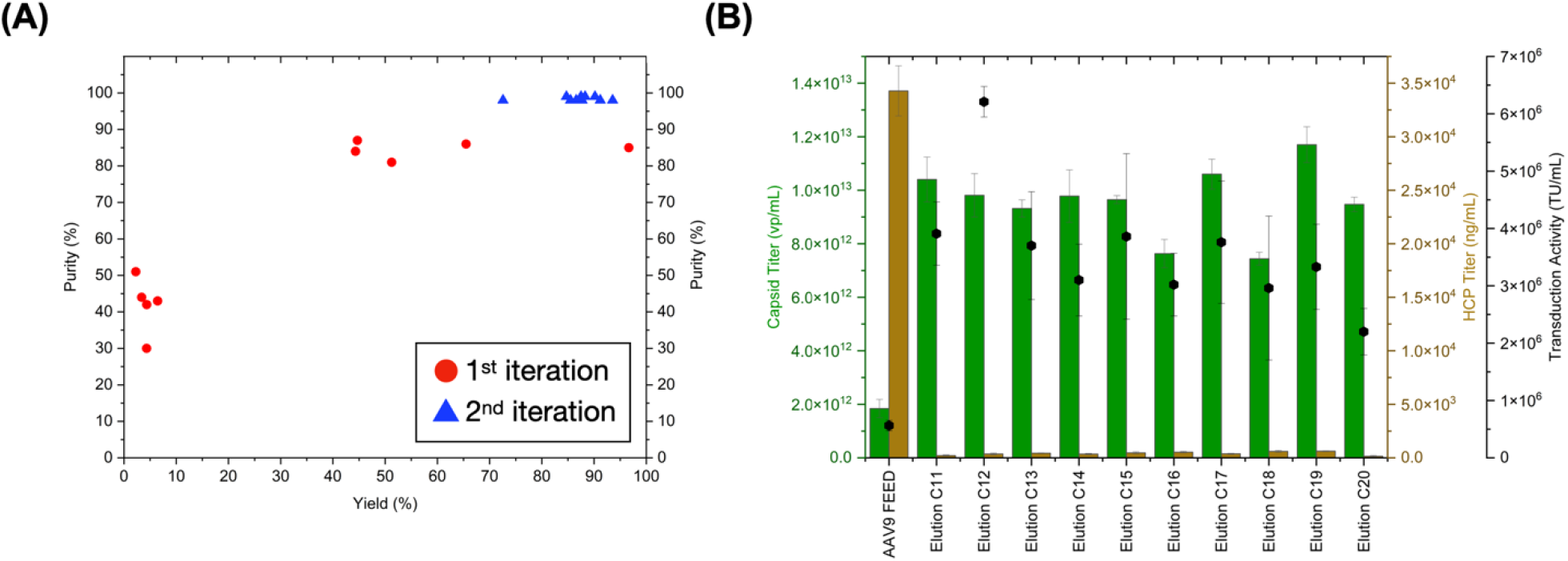
**(A)** Values of capsid yield and purity measured by SEC-HPLC analysis of the elution fractions obtained by purifying AAV9 from HEK293 cell lysates using AAVidity operated with process input parameters selected across three iterations of the closed-loop Bayesian optimization process. **(B)** Values of capsid titer, HEK293 HCP titer, and HT1080 cell transduction activity in the elution fractions obtained by purifying AAV9 using AAVidity resin operated with process parameters selected in the third iteration of Bayesian optimization.

Selected eluates were evaluated by AAV9 capsid ELISA and HEK293 HCP ELISA to quantify capsid recovery and impurity clearance, as well as by HT1080 cell transduction assays via flow cytometry to assess the bioactivity of the purified vectors (**Figure 5B**). Under Bayesian-optimized conditions, the AAVidity resin achieved high capsid titers (7.4·10^12^ – 1.2·10^12^ vp/mL, corresponding to a ∼5.2-fold enrichment) and residual HCP concentrations reaching as little as 146 ng/mL (HCP LRV ∼ 3.15), corresponding to 2.1 μg of HCPs per equivalent Zolgensma dose (∼1.1·10^14^ vp). While above the limit admitted for *in vivo* applications, the value of residual impurities can be easily reduced to meet clinical standards during the post-affinity polishing step^80,81^. Finally, the flow cytometric analysis of AAV9 on HT1080 cells confirmed that the vectors purified under optimized conditions retained high transduction activity, a prerequisite for therapeutic efficacy and safety.

#### 2.3.3. Purification of AAV5

This vector is a genetically distinct member of the AAV family, characterized by unique capsid structural features that influence its cell binding and transduction properties. AAV5 demonstrated efficient transduction of airway epithelial cells, vascular endothelial and smooth muscle cells, neuronal tissues, and hepatocytes, thus emerging a valuable vector for targeting multiple organ systems. Several AAV5-based gene therapies are actively undergoing clinical trials, addressing conditions such as Fabry disease as well as hematology and rare genetic diseases. Two FDA-approved gene therapies utilize AAV5 vectors, namely Roctavian (approved in 2023) and Hemgenix (2022) for the treatment of hemophilia A and hemophilia B, respectively^82–84^. AV5 is the most sequence-divergent and structurally distinct serotype, particularly when compared to AAV2 and AAV9^61,85,86^. Its capsid architecture features unique variations in surface-exposed loops and viral protein domains, which modulate receptor interactions, tissue tropism, and antigenicity. The mechanism of AAV5 capture b AAVidity differs from that of AAV2 and AAV9 binding. The peptide ligand, engineered by hybridizing the AAV5-targeting PKD1 domain and the AAV2/9-targeting PKD2 domain of the AAV receptor, binds AAV2 and AAV9 at physiological pH and releases them under mildly acidic pH, whereas it binds AAV5 under acidic pH and releases it as the pH is increased in presence of magnesium chloride^46,87^. We therefore adjusted the space of input parameters explored by the Bayesian optimization and initialized the first chromatographic test set via random sampling. Notwithstanding the unique AAV5-AAVidity binding mechanism, several conditions were identified within the first two test sets that afforded high product yield (> 94%) and purity (> 99%), as reported in **Figure S5** and **Table S8**. These, however, spanned a broad range of formulations of the wash buffer (1 – 24 mS/cm and pH 4.5 – 5.6) and elution buffer (8.4 – 19.4 mS/cm and pH 6.1 – 8.5). The identification of multiple optima required the design of a third test set to achieve consistently high yield and purity (**Figure 6A**). The quality of the purified AAV5 were confirmed by ELISA, which indicated a capsid titer of 2.6·10^11^ – 3.2·10^12^ vp/mL and residual HCP titer below the ELISA limit of detection (**Figure 6B**). Finally, the transduction assays in HT1080 cells confirmed that, across many diverse purification conditions, the Bayesian-guided purification strategy preserved the functional activity of the isolated AAV5 vectors.

**Figure 6.**
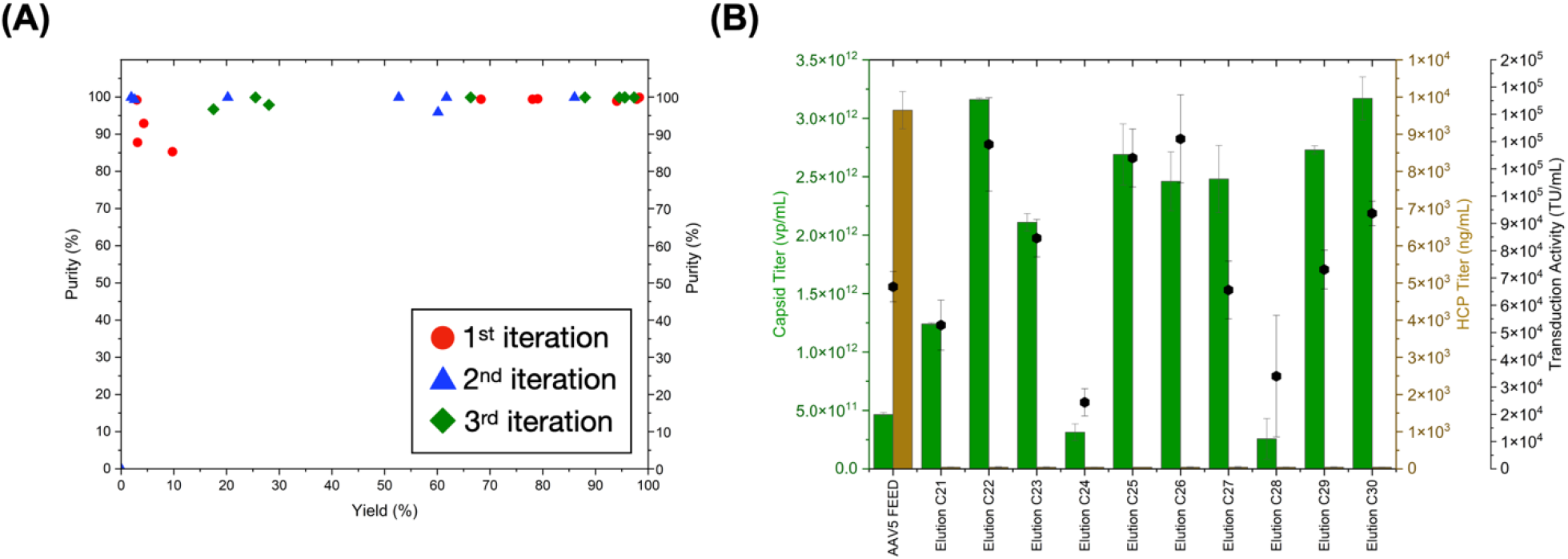
**(A)** Values of capsid yield and purity measured by SEC-HPLC analysis of the elution fractions obtained by purifying AAV5 from HEK293 cell lysates using AAVidity operated with process input parameters selected across three iterations of the closed-loop Bayesian optimization process. **(B)** Values of capsid titer, HEK293 HCP titer, and HT1080 cell transduction activity in the elution fractions obtained by purifying AAV2 using AAVidity resin operated with process parameters selected in the third iteration of Bayesian optimization.

Collectively, these results demonstrate the effectiveness of our closed-loop BO approach for the rapid optimization of purification operations in the gene therapy industry. Especially notable was the observation that, whilst product yield was adopted as the sole optimization objective, the selected input process parameters satisfy other critical quality attributes of viral vectors, chiefly cell transduction activity and purity, thus improving at once therapeutic efficacy, safety, and affordability of gene therapies.

### 2.4. Evaluating the effect of process parameters on optimization objectives

The complexity of gene therapy purification, which stems from the unique design of the viral vector, the variability in the feedstocks, and the wide space of operation parameters of affinity chromatography, makes it challenging to elucidate the relationships between process parameters and the resulting productivity and quality of the isolated viruses. To determine the impact of individual process parameters on model predictions, we analyzed the chromatographic test sets using SHAP (SHapley Additive exPlanations, **Figure 7**)^88,89^. The SHAP analysis demonstrated that the affinity adsorbent AAVidity is ideally suited for the industrial-scale gene therapy biomanufacturing. Firstly, product yield is enhanced by loading large feedstock volumes and performing washing and elution at high flow rates, which reduce respectively the number and the duration of chromatographic campaigns, thereby decreasing the total process time. Secondly, product yield is optimal when employing physiological pH and conductivity in wash buffers, which preserve the vectors’ cell transduction activity, ultimately promoting the efficacy and safety of gene therapies. In this regard, the SHAP analysis complements the quantitative evaluations of process productivity and product quality with a qualitative evaluation of the aptness of a process technology to optimize its operation under conditions that support the stability, integrity, and bioactivity of the product.

**Figure 7.**
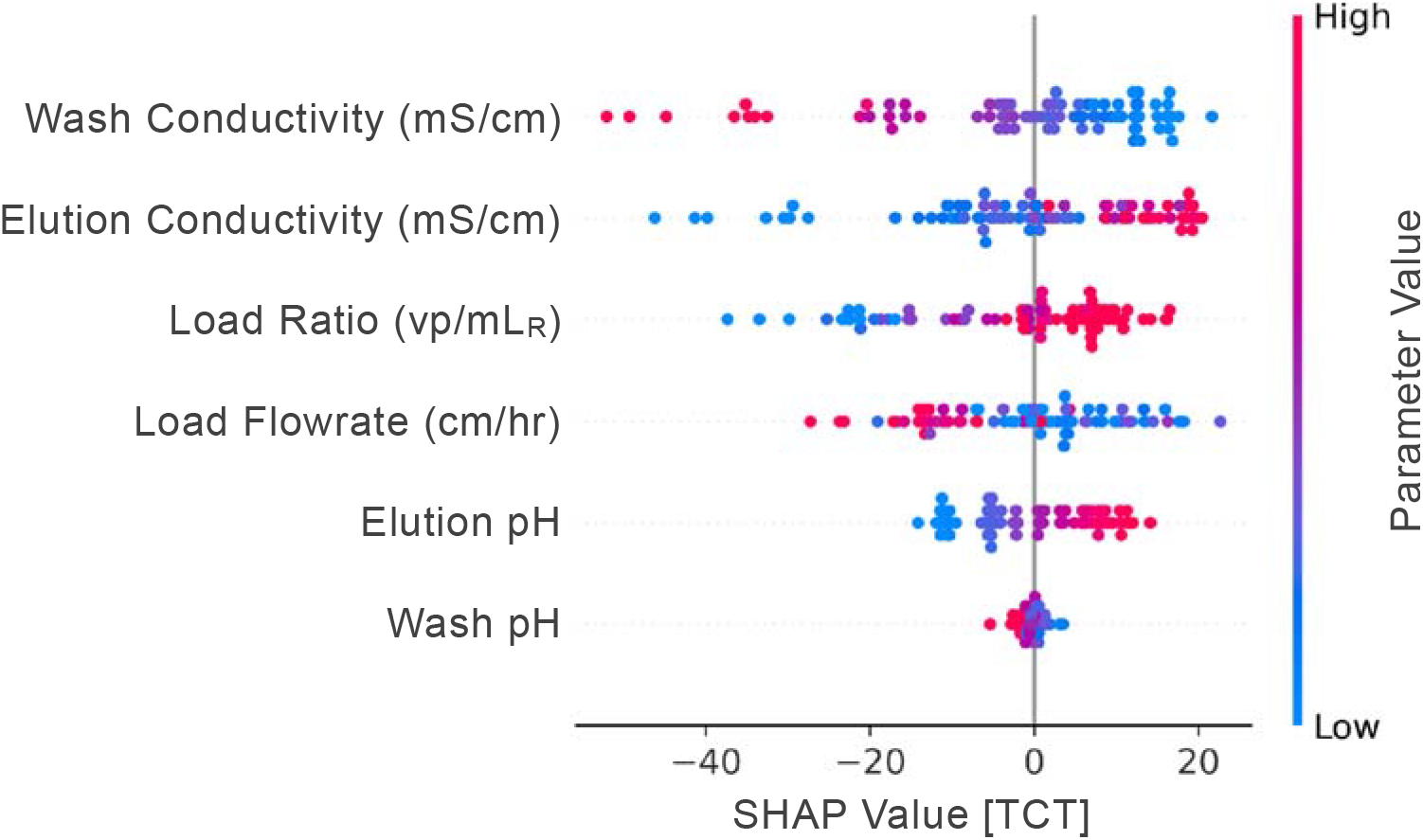
SHAP values illustrate the individual parameter contributions to the total capsid yield obtained by purifying AAV2, AAV5, and AAV9 from HEK293 cell lysates using the affinity resin AAVidity. SHAP analysis quantifies the effect of each chromatographic process parameter, supporting data-driven optimization of process conditions. Red dots represent high parameter values, while blue dots indicate low values. The X-axis displays the magnitude and direction of each parameter’s impact on total capsid production.

## 3. Conclusions

This study demonstrates the effectiveness and versatility of a closed-loop BO workflow for the rapid development of AAV purification processes by affinity chromatography. We leveraged a GP surrogate model trained on extensive historical data to navigate multidimensional space of process design, achieving optimal conditions for three clinically relevant AAV serotypes (AAV2, AAV5, and AAV9) in less than 30 experimental iterations per serotype. The adaptive optimization framework, guided by the *qEI* acquisition function, consistently outperformed random sampling and conventional development approaches, as demonstrated by both simulation and experimental campaigns, achieving a remarkable improvement in yield from the original baseline of 70% to 99%, with a concurrent increase in purity. Our results highlight the product-agnostic applicability of the approach, with rapid transfer of process knowledge between serotypes, despite their unique biomolecular features and distinct binding mechanisms to the affinity resin. Notably, the single-objective optimization for yield also delivered significant improvements in purity and vector bioactivity, indicating a synergistic relationship between these critical quality attributes. Finally, the SHAP analysis further elucidated the influence of key process parameters, revealing that high yields are achieved under conditions that preserve the vector’s bioactivity and process feasibility. The closed-loop, data-driven methodology proposed in this study showed true promise to accelerate process development, reduce experimental burden, and support the production of safer and more efficacious gene therapy vectors. We expect this approach to be readily extensible to other viral vectors and bioprocessing modalities, since its integration with high-throughput analytics and explainable AI tools (*e.g*., SHAP) provides both quantitative and qualitative insights into process performance. In summary, this work establishes a new paradigm for the rational, efficient, and scalable optimization of viral vector purification, offering a path toward more affordable and accessible gene therapies. Future efforts will focus on expanding the framework to other viral vector families (*e.g*., lentivirus and adenovirus), exploring multi-objective optimization, and integrating real-time process analytics for fully autonomous bioprocessing.

## 4. Materials and Methods

### 4.1. Data extraction

All affinity chromatography data were extracted from the UNICORN software integral to the AKTA purification systems. The extracted files were converted using the PyCORN Python package to obtain a non-proprietary data format. The Pandas Python module was used for further preprocessing to compile experimental parameters and values of interest.

### 4.3. Gaussian process modeling

GP was used to predict the total yield of AAV capsids by modeling the experimental space of input parameters defined in **Table S1**. The GP was first implemented and benchmarked using the scikit-learn Python library and was subsequently reimplemented using GPyTorch for compatibility with the Bayesian backend (BayBe) Python package. The kernel used for predicting the “total capsid” yield (*K*_*TC*_) in both implementations is described in **Equations 1 - 4**.

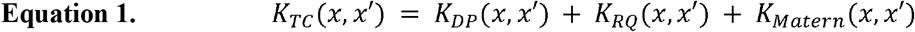

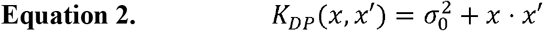

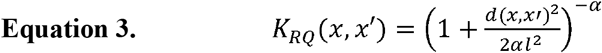

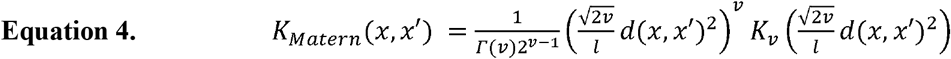

Wherein *K*_*DP*_ is the dot product kernel, *K*_*RQ*_ is the rational quadratic kernel, and *K*_*Matern*_ is the Martern kernel. An expansion of the above equations with defined parameters is described in the Supplementary Information *Section S.1*. Combining these kernels enables the model to capture various functional forms, from linear trends to more complex, smooth, or non-smooth patterns, with different characteristic length scales. The additive combination accounts for that kernel’s contribution, enabling the model to automatically adjust how it weighs each type of relationship based on the input parameters of the process.

### 4.3. Simulation and Evaluation of Acquisition Functions

A simulation of the optimization campaign was conducted to evaluate the performance of four acquisition functions, namely *qExpectedImprovement* (*qEI*; additional details about the *qEI* acquisition function are in the Supplementary Information *Section S.2*), *qLogExpectedImprovement* (*qLogEI*), *qProbabilityofImprovement* (*qPI*), and *qUpperConfidence-Bounds* (*qUCB*). Random sampling provided a baseline policy and was performed by uniformly acquiring parameter sets across the space of process parameters (**Table S1**) to identify an array of input parameters (*x’*) that is expected to maximize the objective function ln[TCT(*x*)] (**Equation 5**). All acquisition functions leveraged quasi-Monte Carlo (qMC) sampling techniques to evaluate the objective function’s integral over the posterior distribution.^28^ To mimic a realistic experimental space, we utilized a model trained on experimental data as a source of ground truth for the simulation.

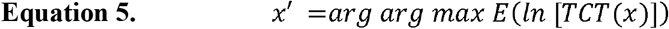

### 4.4. Materials

Phosphate-buffered saline (PBS) tablets, hydrochloric acid, Bis-Tris, magnesium chloride, phosphoric acid, and sodium chloride were purchased from Fisher Chemicals (Hampton, NH, USA). Toyopearl NH□-750F resin was obtained from Tosoh Bioscience (Tokyo, Japan). Gibco™ Viral Production Cells prototype 2.0, AAV-MAX Enhancer, Viral□Plex™ Complexation Buffer, Transfection Reagent, Dulbecco’s Modified Eagle Medium (DMEM), fetal bovine serum (FBS), and AAV-MAX lysis buffer were acquired from ThermoFisher Scientific (Waltham, MA, USA). Plasmids pAAV2/2 (Addgene plasmid #104963), pAAV2/5 (Addgene plasmid #104964), pAAV2/9n (Addgene plasmid #112865), pDGM6 (Addgene plasmid #110660) were gifted by James M. Wilson and David Russell, while plasmid pAAV CAGG eGFP (Addgene plasmid #107707) was a gift from Troy Margrie. Benzonase endonuclease (GENIUS™ Nuclease, 500 KU) was sourced from Santa Cruz Biotechnology (Dallas, TX, USA). The Bis-Tris HCl, magnesium chloride (MgCl_2_), methanol, phosphoric acid, sodium chloride (NaCl), sodium hydroxide (NaOH), Pluronic™ F-68, and potassium chloride (KCl), were purchased from Fisher Chemical (Hampton, NH, USA). The HT1080 cells were from ATCC (Manassas, VA). Aurora Pro Scientific 1□mL FPLC columns were obtained from Fisher Scientific. A BioResolve SEC mAb Column (bead diameter: 2.5 μm; pore diameter: 200Å; column diameter: 7.8 mm; column length: 300 mm) was sourced from Waters Inc. (Milford, MA, USA). The serotype-specific ELISA kits for the quantification of AAV2, AAV5, and AAV9 were purchased from Progen (Heidelberg, Germany), while HEK293 host cell protein (HCP) ELISA kits were acquired from Cygnus Technologies (Southport, NC, USA). The AAVidity affinity resin was acquired from LigaTrap Technologies (Raleigh, NC). The BioResolve SEC mAb column (200□Å, 2.5□μm, 7.8×300□mm) for size exclusion chromatography was obtained from Waters Corporation (Milford, MA, USA).

### 4.5. Production of AAV Feed Material

HEK 293 cells (Gibco™ Viral Production Cells 2.0) were inoculated at a density of 3·10^6^ cells/mL in a culture medium supplemented with 1% v/v AAV-MAX Enhancer. The cultures were maintained under controlled conditions (37°C, 8% CO□, 120 rpm orbital agitation) in a humidified incubator to promote optimal cell expansion. Plasmid constructs comprising pRC2 (encoding AAV Rep and Cap genes), pHelper (providing adenoviral helper functions essential for AAV replication), and pAAV-GFP (harboring the GFP transgene cassette) were co-formulated at the molar ratios listed in **Table 2**, yielding a total nucleic acid concentration of 2 μg/mL relative to the final culture volume. The plasmid mixture was diluted to 10% v/v with Viral Plex™ Complexation Buffer to facilitate complex formation. In parallel, volumes of AAV□MAX Transfection Booster and AAV□MAX Transfection Reagent corresponding to 0.3% v/v and 0.6% v/v of the total culture volume were added and mixed at ambient temperature for 30 minutes maximum to allow for the polyplex formation. The resultant transfection complex was introduced to the cell suspension and incubated for 72 hours (37°C, 8% CO_2_, 120 rpm orbital agitation). Cell lysis was then conducted as described in prior work^47^. The crude lysate was treated with 50 U/mL of GENIUS™ Nuclease (benzonase endonuclease) in the presence of 2 mM MgCl_2_ for 2 hours at 37°C to degrade host and plasmid DNA. The lysates were clarified by centrifugation at 4500×g for 30 minutes, after which the clarified supernatants were stored at -80□ before the affinity purification.

**Table 2.** Plasmid ratios for the production of different AAV serotypes.

| Serotype | Transfection | Plasmids Ratio |
| --- | --- | --- |
|  |  | (pRC : pHelper: pAAV-GFP or pRC+Helper : pAAV-GFP) |
| AAV2 | Triple | 2:0.5:1 |
| AAV5 | Triple | 2:0.5:1 |
| AAV6 | Double | 1:1 |
| AAV9 | Triple | 2:0.5:1 |

### 4.6. AAV purification by affinity chromatography

The buffer composition of the clarified HEK293 cell lysates containing AAV2 and AAV9 was initially adjusted to a final NaCl concentration of 150 mM and a pH of 7.0, while the lysate containing AAV5 was adjusted to 150 mM NaCl and pH 5.0. The feedstocks were then concentrated and diafiltered by tangential flow filtration (TFF) using Pellicon® 2 Cassette Ultrafiltration Biomax^®^ Module (MWCO ∼100kDa; Millipore Sigma, Darmstadt, Germany): the feedstocks containing AAV2 and AAV9 were conditioned to 20 mM NaCl and 2 mM MgCl2 in 10 mM Bis-Tris at pH 7.0, while the feedstocks containing AAV5 were conditioned to 2 mM MgCl2 in 50 mM sodium acetate at pH 5.0. The purification of all AAVs by affinity chromatography was conducted on an AKTA Avant 25 FPLC system (Cytiva, Uppsala, Sweden) using 1 mL column (diameter: 0.5 cm; height: 5cm) was packed with AAVidity resin. The resin was equilibrated with 20 column volumes (CVs) of 20 mM NaCl and 2 mM MgCl2 in 10 mM Bis-Tris at pH 7.0 before loading the AAV2 and AAV9 feedstocks, or 2 mM MgCl2 in 50 mM sodium acetate at pH 5.0 before loading the AAV5 feedstock. The buffer composition of the washing and elution steps (10 and 5 CVs, respectively) was adjusted as listed in **Tables S3 – S5**. After elution, the resin was sanitized with 0.1 M NaOH (aq.) for a total contact time of 30 mins, washed with PBS at pH 2.0, and equilibrated with 20 mM NaCl and 2 mM MgCl2 in 10 mM Bis-Tris at pH 7.0. The column was stored in 20% v/v ethanol. All chromatographic steps except the loading step were performed at 1 mL/min (306 cm/hr).

### 4.7. Analysis of process samples by size-exclusion analytical chromatography (SEC-HPLC)

The process samples including the clarified feedstocks and the flow-through, wash, and elution fractions were analyzed by SEC-HPLC using a BioResolve SEC column (Waters, Milford, MA). The analysis was conducted on a sample volume of 10 μL using a 40-minute isocratic method using 0.2 M potassium chloride and 0.05% w/w sodium azide in 50 mM sodium phosphate buffer at pH 7.0 and the flow rate of 0.50 mL/min. The SEC column effluent was continuously monitored by UV spectroscopy (260 and 280 nm) and fluorescence spectroscopy (exc/em: 280/350 nm). The analysis of the SEC chromatograms – namely signal normalization, baseline identification, peak detection, and area integration – was performed using the Scipy Python package. To ensure proper peak detection, all data was processed using the Savitzky-Golay smoothing function, with a window length of 11 and a polynomial order of 0. The monomeric AAV purity was estimated as the ratio between the area of the peak assigned to the AAV capsids (retention time, RT: 10 - 11 min) and the total area of the peaks associated with impurities (RT: 8 - 10 min and 11 - 20 min, **Figure S6**).

### 4.8. Quantification of AAV capsid titer and HEK293 HCP titer by ELISA

The titers of HEK293 host cell proteins (HCPs, μg/mL) and AAV capsids (vp/mL) in the clarified feedstocks and in the affinity eluates were measured using Generation 3 HEK293 HCP ELISA kit (Cygnus Technologies, Southport, NC) and serotype-specific AAV ELISA kits (Progen, Wayne, PA), respectively.

### 4.9. Transduction Assay

The HT1080 cells were cultured in DMEM medium containing 10% v/v FBS at 37°C in a 5% CO_2_ environment until reaching 80% confluence. Thereafter, the cells were seeded into 96-well plates at the density of 10,000 cells per well and incubated overnight. The clarified feedstocks and affinity eluates containing AAVs were serially diluted in DMEM. A 0.1 mL volume of each diluted sample was added to the HT1080 cells. After 24 hours, the medium was replaced with fresh DMEM containing 10% v/v FBS, and the cells were cultured for an additional 24 hours. The quantification of green fluorescent protein (GFP^+^) expression was performed using a CytoFlex flow cytometer (Beckman Coulter, Brea, CA). The transduction units per milliliter (TU/mL) were subsequently calculated based on the established **Equation 6**, wherein N_HT1080_ is the number of cells incubated with the diluted AAV sample, V is the volume of the diluted AAV sample, and DF is the dilution factor.

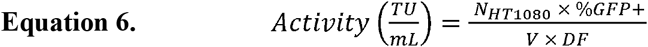

## Data availability

All data supporting the findings and conclusions presented in this study are contained within the main text and the supplementary information.

## Code availability

All computational methods were created using open-source Python libraries, as comprehensively listed in the supplementary materials. Our code files are available at https://github.com/idanwekhai/VVIRAL.

## Acknowledgments

KPI would like to thank Sobhana A. Sripada for the insightful discussions. All authors wish to acknowledge the funding provided by the Food and Drug Administration (Grant R01FD007481), the Novo Nordisk Foundation (AIM-Bio Grant NNF19SA0035474), the Triangle Universities Center for Advanced Studies Inc. (TUCASI), and the North Carolina Viral Vector Initiative in Research and Learning (NC-VVIRAL) at NC State University.

## Contributions

KPI wrote software to curate data, train models, and guide experiments; SA, AM, MH, and EB conducted all chromatographic and analytical work, and contributed to manuscript composition; ENM, MAD, SM, and AT coordinated the project and contributed to manuscript composition and editing.

