## Supplementary Information for "Adaptive Machine Learning Framework enables Unprecedented Yield and Purity of Adeno-Associated Viral Vectors for Gene Therapy"

**S.1. Gaussian Process Kernel.** The Gaussian process model employs a composite kernel structure formed by the additive composition of three functions, namely the Dot Product kernel (*K_DP_*), the Rational Quadratic kernel (*K_RQ_*), and the Matern kernel (*K_Matern_*). The additive composition of the three kernels yields a flexible covariance structure capable of modeling complex functions with varying degrees of smoothness, multiple characteristic length scales, and both stationary and non-stationary components. The composite kernel (*K_C_*) is expressed by **Equation S1**:

**Equation S1.**	*K_C_ (x,x')* = *K_DP_ (x, x')* + *K_RQ_ (x, x')* + *K_Matern_ (x, x')*

Wherein each component contributes unique properties to the overall covariance structure, enabling the model to capture different aspects of the underlying function space.

Specifically, the Dot Product kernel (*K_DP_*) introduces linear relationships between input features and is defined by **Equation S2**:

**Equation S2.**	$K_{DP}(x, x')=\sigma_{0}^{2}+x\cdot x'$

Wherein $x\cdot x'$ denotes the dot product of input vectors $x$ and $x'$, and $\sigma_{0}^{2}$ is a hyperparameter that controls the kernel’s inhomogeneity. This kernel becomes equivalent to a linear kernel when $\sigma_{0}^{2}=0$. The presence of $\sigma_{0}^{2}$ allows the kernel to model non-stationary functions where the covariance between points depends on their absolute locations in the input parameter space, not just their relative distances. For our implementation, we set $\sigma_{0}^{2}= 0.01$, introducing a small degree of inhomogeneity while maintaining predominantly linear correlation patterns.

The Rational Quadratic kernel (*K_RQ_*) can be interpreted as a scale mixture of Squared Exponential kernels with different length scales. *K_RQ_* is defined by **Equation S3**:

**Equation S3.**	${K_{RQ}(x, x')=\left( 1+\frac{{d(x, x')}^{2}}{2\alpha l^{2}} \right)}^{-\alpha}$

Wherein $d(x, x')$ represents the Euclidean distance between input vectors $x$ and $x'$, defined as $d(x, x') = ||x - x'||$, $\alpha> 0$ is the scale mixture parameter that determines the relative weighting of large-scale and small-scale variations, and $l>0$ is the length scale parameter that governs the characteristic distance over which function values become uncorrelated. The Rational Quadratic kernel approaches the Squared Exponential kernel as $\alpha\to\infty$. However, with finite α values, it can accommodate functions that vary over multiple length scales. In our implementation, we set $\alpha=0.01$ and $l=0.01$, which configures the kernel to model functions with relatively abrupt changes while maintaining sensitivity to fine-grained patterns in the data.

Finally, the Matern kernel introduces controlled function smoothness and is particularly useful for modeling physical processes. *K_Matern_* is defined by **Equation S4**:

**Equation S4.**	$K_{Matern}(x, x') =\frac{1}{\Gamma(v)2^{v-1}}\left( \frac{\sqrt{2v}}{l}{d(x, x')}^{2} \right)^{v}K_{v}\left( \frac{\sqrt{2v}}{l}{d(x, x')}^{2} \right)$

Wherein, $d(x, x')$ represents the Euclidean distance between input vectors $x$ and $x'$, $K_{v}(.)$ is the modified Bessel function of the second kind of order $v$, $\Gamma(.)$ is the gamma function, $v>0$ is the smoothness parameter that controls the differentiability of the resulting function, and $l>0$ is the length scale parameter. The parameter $v$ directly determines the mean-square differentiability of the resulting Gaussian process. Specifically, a process with a Matern covariance function is [$v$]-times differentiable in the mean-square sense. As $v\to\infty$, the Matern kernel converges to the Squared Exponential kernel, resulting in infinitely differentiable functions. In our implementation, we set $l=0.1$ and $v=1.5$, the resulting process is once differentiable, making it suitable for modeling physical phenomena that exhibit some smoothness but are not infinitely differentiable. The selected length scale of $l=0.1$ allows the model to capture medium-scale variations in the data.

**S.2. Acquisition Functions.** The *qExpectedImprovement* (*qEI*) acquisition function is an extension of the standard Expected Improvement (EI) used in Bayesian optimization and is designed for batch optimization scenarios. It quantifies the expected improvement over the current best-observed value when evaluating q candidate points simultaneously. By considering joint information gain across multiple evaluations, *qEI* helps efficiently explore and exploit the search space, making it particularly useful for parallelized optimization settings. It is commonly employed in applications where multiple experiments or simulations can be run concurrently, enhancing optimization efficiency. The *qExpectedImprovement* acquisition function is expressed by **Equation S5**:

**Equation S5.**	$qEI(X) = E_{Y\sim p(Y|X, D)}[max(0, max(Y) - f(X^{+}))]$

Wherein *X = {x_1_, x_2_, …, x_q_}* is the batch of q candidates points being evaluated, *Y = {y_1_, y_2_, …, y_q_}* represents the corresponding function values of these points, *p(Y|X, D)* is the predictive distribution given by the Gaussian process model, conditioned on the observed data *D,* $f(X^{+})$ is the current best observed function value in the dataset, and $E$ denotes the expected value operator.

Due to the complex nature of the joint distribution over multiple points, qEI is typically computed using the Monte Carlo estimation in **Equation S6**:

**Equation S6.**	$qEI(X) \approx\frac{1}{N}\sum_{i=1}^{N} max(0, max(Y^{(i)}) - f(X^{+}))$

Wherein $Y^{(i)}\sim p(Y|X, D)$ represents the *i-th* sample drawn from the predictive distribution, and *N* the number of Monte Carlo samples. The qEI acquisition function inherently balances exploration (uncertainty sampling) and exploitation (improvement targeting).

***Table S1.*** *Chromatographic process parameters used for the closed-loop Bayesian optimization of the purification of AAV2 and AAV9 from HEK293 cell lysates using AAVidity resin.*

| **Parameter Name** | **Type** | **Range** |
| --- | --- | --- |
| Pure sample (True / False) | Discrete (fixed) | False |
| Serotype | Categorical | AAV2, AAV9 |
| Affinity resin | Categorical (fixed) | AAVidity |
| Elution pH | Discrete Numerical | 5.0 - 9.0, 0.5 step size |
| Wash pH | Discrete Numerical | 5.0 - 9.0, 0.5 step size |
| Equilibration pH | Discrete (fixed) | 7.0 |
| Elution Conductivity (mS/cm) | Continuous | 10.0 - 100.0 |
| Wash Conductivity (mS/cm) | Discrete Numerical | 1.0 - 15.0, 1.0 step size |
| Equilibration Conductivity | Discrete (fixed) | 2.5 |
| Sample Volume (mL) | Discrete Numerical | 5, 10, 15, 20, 25, 30 |
| System Flowrate (cm/h) | Discrete (fixed) | 306 |
| Sample Flowrate (cm/h) | Continuous | 130, 600 |
| From (LFT/ELU) | Discrete | ELU |

***Table S2.*** *Chromatographic process parameters used for the closed-loop Bayesian optimization of the purification of AAV5 from HEK293 cell lysates using AAVidity resin.*

| **Parameter Name** | **Type** | **Range** |
| --- | --- | --- |
| Pure sample (True / False) | Discrete (fixed) | False |
| Serotype | Categorical | AAV5 |
| Affinity resin | Categorical (fixed) | AAVidity |
| Elution pH | Discrete Numerical | 6.0 - 9.0, 0.5 step size |
| Wash pH | Discrete Numerical | 6.0 - 9.0, 0.5 step size |
| Equilibration pH | Discrete (fixed) | 7.0 |
| Elution Conductivity (mS/cm) | Continuous | 0.0 - 20.0 |
| Wash Conductivity (mS/cm) | Discrete Numerical | 0.0 - 10.0, 1.0 step size |
| Equilibration Conductivity | Discrete (fixed) | 2.5 |
| Sample Volume (mL) | Discrete Numerical | 5, 10, 15, 20, 25, 30 |
| System Flowrate (cm/h) | Discrete (fixed) | 306 |
| Sample Flowrate (cm/h) | Continuous | 130, 600 |
| From (LFT/ELU) | Discrete | ELU |

***Table S3.*** *Target and actual conditions utilized for purifying AAV2 from HEK293 cell lysates using AAVidity resin by three iterations of Bayesian optimization.*

| **Test #** | **Wash pH** | | **Wash Conductivity**  **(mS/cm)** | | **Elution pH** | | **Elution Conductivity**  **(mS/cm)** | | **Load Volume**  **(mL)** | **Load Flowrate**  **(cm/hr)** |
| --- | --- | --- | --- | --- | --- | --- | --- | --- | --- | --- |
|  | **Target** | **Actual** | **Target** | **Actual** | **Target** | **Actual** | **Target** | **Actual** |  |  |
| B1R1 | 8.5 | 8.55 | 5 | 6.27 | 7.0 | 7.08 | 92.1 | 84.28 | 5 | 102 |
| B1R2 | 7.5 | 7.57 | 15 | 17.71 | 6.5 | 6.56 | 17.8 | 11.47 | 30 | 102 |
| B1R3 | 6.5 | 6.53 | 10 | 12.23 | 8.5 | 8.45 | 68.7 | 56.18 | 20 | 102 |
| B1R4 | 7.0 | 7.05 | 2 | 3.11 | 8.5 | 8.47 | 99.5 | 95.73 | 15 | 102 |
| B1R5 | 5.0 | 5.05 | 13 | 16.18 | 6.0 | 6.03 | 58.7 | 46.17 | 30 | 102 |
| B1R6 | 6.5 | 6.55 | 1 | 1.70 | 8.0 | 8.05 | 93.5 | 86.89 | 10 | 102 |
| B1R7 | 7.0 | 6.99 | 3 | 3.82 | 5.0 | 5.06 | 32.8 | 23.71 | 30 | 102 |
| B1R8 | 9.0 | 9.06 | 1 | 1.40 | 6.0 | 6.07 | 75.9 | 62.98 | 15 | 102 |
| B1R9 | 6.0 | 6.02 | 13 | 16.02 | 9.0 | 9.02 | 37.5 | 27.59 | 5 | 102 |
| B1R10 | 5.5 | 5.56 | 5 | 6.72 | 7.5 | 7.48 | 35.2 | 25.45 | 5 | 102 |
| B2R1 | 6.5 | 6.51 | 4 | 5.90 | 5.0 | 4.97 | 39.5 | 27.42 | 30 | 175 |
| B2R2 | 6.5 | 6.52 | 11 | 13.9 | 8.5 | 8.41 | 41.8 | 30.15 | 20 | 187 |
| B2R3 | 7.0 | 7.06 | 4 | 5.71 | 5.0 | 5.07 | 49.3 | 36.04 | 25 | 167 |
| B2R4 | 7.0 | 6.95 | 3 | 4.07 | 8.5 | 8.20 | 53.1 | 40.02 | 15 | 173 |
| B2R5 | 7.0 | 6.95 | 3 | 5.61 | 5.0 | 5.07 | 33.7 | 22.24 | 30 | 139 |
| B2R6 | 5.0 | 5.09 | 15 | 19.89 | 6.0 | 6.10 | 27.2 | 17.97 | 30 | 155 |
| B2R7 | 5.0 | 5.05 | 15 | 18.84 | 6.0 | 6.08 | 33.2 | 22.51 | 30 | 160 |
| B2R8 | 9.0 | 8.91 | 1 | 1.19 | 6.0 | 6.06 | 36.5 | 25.41 | 15 | 166 |
| B2R9 | 7.0 | 7.05 | 2 | 3.12 | 5.0 | 5.09 | 84.2 | 74.61 | 30 | 151 |
| B2R10 | 7.0 | 7.05 | 3 | 3.98 | 5.0 | 5.02 | 104.2 | 99.28 | 30 | 131 |
| B3R1 | 5.0 | 5.06 | 4 | 5.11 | 5.0 | 5.07 | 65.3 | 61.04 | 30 | 252 |
| B3R2 | 7.0 | 6.95 | 10 | 11.17 | 6.0 | 5.99 | 69.7 | 64.92 | 30 | 249 |
| B3R3 | 6.5 | 6.49 | 6 | 7.10 | 6.5 | 6.55 | 62.5 | 56.97 | 30 | 406 |
| B3R4 | 6.5 | 6.52 | 7 | 8.01 | 7.0 | 7.03 | 67.2 | 62.53 | 30 | 130 |
| B3R5 | 5.5 | 5.54 | 5 | 6.35 | 7.5 | 7.49 | 81.0 | 81.71 | 30 | 389 |
| B3R6 | 7.0 | 6.97 | 7 | 7.80 | 7.5 | 7.49 | 48.9 | 41.93 | 30 | 453 |
| B3R7 | 7.5 | 7.49 | 2 | 2.44 | 7.5 | 7.46 | 85.1 | 86.72 | 30 | 228 |
| B3R8 | 6.0 | 6.03 | 1 | 2.06 | 8.0 | 7.99 | 51.5 | 45.08 | 30 | 130 |
| B3R9 | 6.0 | 6.01 | 4 | 5.23 | 8.0 | 8.05 | 73.7 | 71.69 | 30 | 231 |
| B3R10 | 8.0 | 8.10 | 5 | 5.69 | 8.0 | 7.95 | 67.7 | 63.43 | 30 | 300 |

***Table S4.*** *Target and actual conditions utilized for purifying AAV9 from HEK293 cell lysates using AAVidity resin by three iterations of Bayesian optimization.*

| **Test #** | **Wash pH** | | **Wash Conductivity**  **(mS/cm)** | | **Elution pH** | | **Elution Conductivity**  **(mS/cm)** | | **Load Volume**  **(mL)** | **Load Flowrate**  **(cm/hr)** |
| --- | --- | --- | --- | --- | --- | --- | --- | --- | --- | --- |
|  | **Target** | **Actual** | **Target** | **Actual** | **Target** | **Actual** | **Target** | **Actual** |  |  |
| B1R1 | 5.0 | 4.95 | 1 | 2.20 | 9.0 | 8.91 | 95.55 | 88.00 | 30 | 544.86 |
| B1R2 | 9.0 | 9.01 | 15 | 15.53 | 5.0 | 4.95 | 10.18 | 15.32 | 5 | 588.67 |
| B1R3 | 5.0 | 5.20 | 15 | 16.54 | 5.0 | 5.12 | 11.11 | 13.73 | 30 | 473.29 |
| B1R4 | 5.0 | 5.10 | 1 | 2.02 | 5.0 | 5.15 | 100.34 | 95.00 | 5 | 594.99 |
| B1R5 | 9.0 | 9.01 | 1 | 1.25 | 9.0 | 9.05 | 10.28 | 14.19 | 15 | 136.63 |
| B1R6 | 5.0 | 5.20 | 15 | 16.32 | 9.0 | 8.91 | 100.62 | 95.10 | 15 | 528.09 |
| B1R7 | 9.0 | 8.70 | 9 | 9.78 | 5.0 | 4.98 | 99.70 | 95.00 | 30 | 227.32 |
| B1R8 | 9.0 | 9.10 | 10 | 10.6 | 9.0 | 8.05 | 11.02 | 14.70 | 10 | 396.14 |
| B1R9 | 5.0 | 5.20 | 1 | 2.14 | 5.0 | 5.10 | 10.44 | 18.04 | 25 | 454.59 |
| B1R10 | 9.0 | 9.10 | 1 | 1.20 | 5.0 | 4.96 | 100.24 | 94.5 | 10 | 586.26 |
| B2R1 | 9.0 | 9.15 | 1 | 1.175 | 9.0 | 9.30 | 10.53 | 14.19 | 30 | 441.89 |
| B2R2 | 9.0 | 9.14 | 1 | 1.15 | 5.0 | 4.96 | 100.22 | 96.10 | 30 | 242.35 |
| B2R3 | 9.0 | 9.11 | 1 | 1.17 | 9.0 | 8.80 | 99.61 | 91.70 | 25 | 563.79 |
| B2R4 | 5.0 | 5.09 | 1 | 2.13 | 9.0 | 8.74 | 99.59 | 93.20 | 20 | 563.27 |
| B2R5 | 9.0 | 8.95 | 1 | 1.20 | 7.0 | 7.14 | 10.61 | 14.60 | 30 | 234.49 |
| B2R6 | 9.0 | 8.90 | 4 | 4.48 | 9.0 | 8.78 | 99.97 | 92.20 | 25 | 559.79 |
| B2R7 | 8.0 | 8.03 | 1 | 1.22 | 8.5 | 8.13 | 10.58 | 14.18 | 30 | 264.91 |
| B2R8 | 9.0 | 8.96 | 1 | 1.15 | 6.5 | 6.58 | 99.91 | 89.00 | 25 | 505.47 |
| B2R9 | 9.0 | 8.93 | 2 | 2.25 | 7.0 | 7.08 | 100.27 | 94.2 | 30 | 269.68 |
| B2R10 | 9.0 | 8.90 | 3 | 3.39 | 5.0 | 5.02 | 10.87 | 15.89 | 30 | 262.04 |

***Table S5.*** *Target and actual conditions utilized for purifying AAV5 from HEK293 cell lysates using AAVidity resin by three iterations of Bayesian optimization.*

| **Test #** | **Wash pH** | | **Wash Conductivity**  **(mS/cm)** | | **Elution pH** | | **Elution Conductivity**  **(mS/cm)** | | **Load Volume**  **(mL)** | **Load Flowrate**  **(cm/hr)** |
| --- | --- | --- | --- | --- | --- | --- | --- | --- | --- | --- |
|  | **Target** | **Actual** | **Target** | **Actual** | **Target** | **Actual** | **Target** | **Actual** |  |  |
| B1R1 | 5.0 | 4.95 | 9.0 | 18.92 | 6.0 | 6.06 | 13.94 | 10.31 | 10 | 382.13 |
| B1R2 | 5.0 | 4.90 | 9.0 | 23.27 | 6.5 | 6.57 | 17.77 | 10.38 | 15 | 463.82 |
| B1R3 | 5.5 | 5.54 | 2.0 | 4.45 | 7.0 | 7.10 | 2.73 | 3.11 | 20 | 148.66 |
| B1R4 | 5.0 | 5.06 | 1.0 | 22.07 | 7.0 | 6.10 | 16.59 | 1.8.0 | 20 | 293.49 |
| B1R5 | 4.0 | 4.06 | 7.0 | 16.74 | 6.5 | 6.45 | 11.95 | 8.50 | 20 | 277.96 |
| B1R6 | 5.0 | 5.05 | 2.0 | 18.76 | 7.5 | 7.55 | 13.93 | 3.11 | 5 | 236.70 |
| B1R7 | 4.5 | 4.55 | 5.0 | 12.31 | 6.0 | 6.06 | 8.38 | 6.30 | 25 | 243.47 |
| B1R8 | 5.5 | 5.40 | 8.0 | 3.58 | 8.0 | 8.08 | 2.21 | 9.42 | 20 | 305.13 |
| B1R9 | 5.5 | 5.50 | 8.0 | 18.87 | 7.0 | 7.03 | 12.39 | 9.32 | 20 | 296.63 |
| B1R10 | 5.5 | 5.56 | 5.0 | 24.08 | 8.0 | 8.09 | 18.27 | 6.27 | 15 | 323.25 |
| B2R1 | 4.0 | 4.10 | 5.0 | 6.09 | 6.0 | 6.10 | 0.00 | 0.344 | 30 | 130.77 |
| B2R2 | 5.5 | 5.39 | 0.0 | 0.80 | 6.0 | 6.00 | 0.89 | 1.754 | 30 | 208.01 |
| B2R3 | 4.0 | 4.08 | 6.0 | 6.72 | 6.0 | 6.01 | 2.29 | 3.78 | 30 | 555.61 |
| B2R4 | 4.0 | 4.06 | 1.0 | 1.88 | 8.0 | 7.94 | 0.08 | 0.20 | 30 | 595.08 |
| B2R5 | 5.5 | 5.50 | 4.0 | 4.98 | 6.0 | 6.01 | 1.01 | 1.94 | 30 | 543.99 |
| B2R6 | 4.0 | 4.07 | 7.0 | 8.2 | 6.0 | 5.93 | 18.73 | 25.06 | 30 | 161.58 |
| B2R7 | 5.5 | 5.42 | 2.0 | 2.97 | 8.5 | 8.44 | 0.14 | 0.361 | 30 | 141.26 |
| B2R8 | 5.5 | 5.41 | 6.0 | 6.99 | 6.0 | 5.93 | 19.56 | 25.64 | 5 | 130.66 |
| B2R9 | 4.0 | 4.09 | 0.0 | 0.83 | 8.5 | 8.54 | 19.39 | 24.00 | 5 | 570.52 |
| B2R10 | 4.0 | 4.08 | 3.0 | 4.00 | 6.0 | 6.04 | 0.034 | 0.313 | 30 | 586.77 |
| B3R1 | 4.0 | 3.95 | 5.0 | 7.01 | 6.0 | 5.95 | 14.86 | 21.56 | 30 | 212.18 |
| B3R2 | 5.5 | 5.51 | 3.0 | 4.83 | 6.0 | 6.04 | 8.12 | 13.27 | 30 | 130 |
| B3R3 | 4.0 | 4.04 | 5.0 | 6.99 | 8.5 | 8.32 | 15.88 | 22.99 | 30 | 600 |
| B3R4 | 4.0 | 4.04 | 5.0 | 6.94 | 8.5 | 8.57 | 11.04 | 16.31 | 5 | 130 |
| B3R5 | 5.5 | 5.43 | 1.0 | 2.31 | 8.5 | 8.58 | 20.00 | 27.59 | 30 | 130 |
| B3R6 | 5.5 | 5.54 | 1.0 | 5.69 | 6.0 | 5.83 | 8.97 | 14.13 | 30 | 130 |
| B3R7 | 4.0 | 3.97 | 4.0 | 4.81 | 6.0 | 6.11 | 16.76 | 24.60 | 30 | 130 |
| B3R8 | 4.0 | 3.96 | 3.0 | 4.74 | 8.5 | 8.41 | 7.68 | 12.46 | 5 | 130 |
| B3R9 | 5.5 | 5.59 | 3.0 | 4.53 | 6.0 | 5.96 | 20.00 | 24.01 | 30 | 600 |
| B3R10 | 5.5 | 5.58 | 0.0 | 0.92 | 8.5 | 8.57 | 16.08 | 23.60 | 30 | 130 |

***Table S6.*** *Values of capsid yield and purity measured by SEC-HPLC analysis of the elution fractions obtained by purifying AAV2 from HEK293 cell lysates using AAVidity operated with process input parameters selected across three iterations of the closed-loop Bayesian optimization process.*

| **Test #** | **Chromatographic Set 1** | | **Chromatographic Set 2** | | **Chromatographic Set 3** | |
| --- | --- | --- | --- | --- | --- | --- |
|  | Yield (%) | Purity(%) | Yield (%) | Purity(%) | Yield (%) | Purity(%) |
| **1** | 45.4 | 97.3 | 92.4 | 97.6 | 85.5 | 97.1 |
| **2** | 4.5 | 89.9 | 75.0 | 97.2 | 84.8 | 97.8 |
| **3** | 55.9 | 99.1 | 91.6 | 95.9 | 92.2 | 95.9 |
| **4** | 60.8 | 94.1 | 87.1 | 95.3 | 92.0 | 96.1 |
| **5** | 24.1 | 99.9 | 82.1 | 95.9 | 94.0 | 95.8 |
| **6** | 53.9 | 85.9 | 5.92 | 93.7 | 99.9 | 97.1 |
| **7** | 56.6 | 92.6 | 5.99 | 98.1 | 97.4 | 94.3 |
| **8** | 58.7 | 97.3 | 75.9 | 93.8 | 97.7 | 95.0 |
| **9** | 22.9 | 89.3 | 75.8 | 90.6 | 99.0 | 94.7 |
| **10** | 44.9 | 96.5 | 75.6 | 89.7 | 96.6 | 94.5 |

***Table S7.*** *Values of capsid yield and purity measured by SEC-HPLC analysis of the elution fractions obtained by purifying AAV9 from HEK293 cell lysates using AAVidity operated with process input parameters selected across three iterations of the closed-loop Bayesian optimization process.*

| **Test #** | **Chromatographic Set 1** | | **Chromatographic Set 2** | |
| --- | --- | --- | --- | --- |
|  | Yield (%) | Purity(%) | Yield (%) | Purity(%) |
| **1** | 96.7 | 84.6 | 87.5 | 98.7 |
| **2** | 2.3 | 51.1 | 93.6 | 98.2 |
| **3** | 4.3 | 29.6 | 91.2 | 98.4 |
| **4** | 44.7 | 86.8 | 87.7 | 98.0 |
| **5** | 65.5 | 86.0 | 90.2 | 98.9 |
| **6** | 4.4 | 42.4 | 72.6 | 98.1 |
| **7** | 6.5 | 42.8 | 88.3 | 98.78 |
| **8** | 3.4 | 44.4 | 86.6 | 98.3 |
| **9** | 44.3 | 84.1 | 85.5 | 98.3 |
| **10** | 51.3 | 81.4 | 84.7 | 98.7 |

***Table S8.*** *Values of capsid yield and purity measured by SEC-HPLC analysis of the elution fractions obtained by purifying AAV5 from HEK293 cell lysates using AAVidity operated with process input parameters selected across three iterations of the closed-loop Bayesian optimization process.*

| **Test #** | **Chromatographic Set 1** | | **Chromatographic Set 2** | | **Chromatographic Set 3** | |
| --- | --- | --- | --- | --- | --- | --- |
|  | Yield (%) | Purity(%) | Yield (%) | Purity(%) | Yield (%) | Purity(%) |
| **1** | 4.3 | 92.9 | 0.0 | 0.0 | 25.94 | 99.9 |
| **2** | 3.1 | 87.8 | 0.0 | 0.0 | 97.34 | 99.9 |
| **3** | 79.1 | 99.5 | 0.0 | 0.0 | 66.33 | 99.9 |
| **4** | 97.8 | 99.4 | 1.9 | 99.9 | 28.04 | 99.9 |
| **5** | 68.3 | 99.4 | 2.5 | 99.3 | 95.61 | 99.9 |
| **6** | 78.1 | 99.4 | 61.8 | 99.9 | 95.57 | 99.9 |
| **7** | 98.3 | 99.9 | 52.7 | 99.9 | 95.50 | 99.9 |
| **8** | 2.9 | 99.2 | 60.1 | 95.9 | 17.53 | 99.9 |
| **9** | 9.7 | 85.3 | 86.0 | 99.9 | 88.01 | 99.9 |
| **10** | 94.1 | 98.9 | 20.2 | 99.9 | 94.54 | 99.9 |

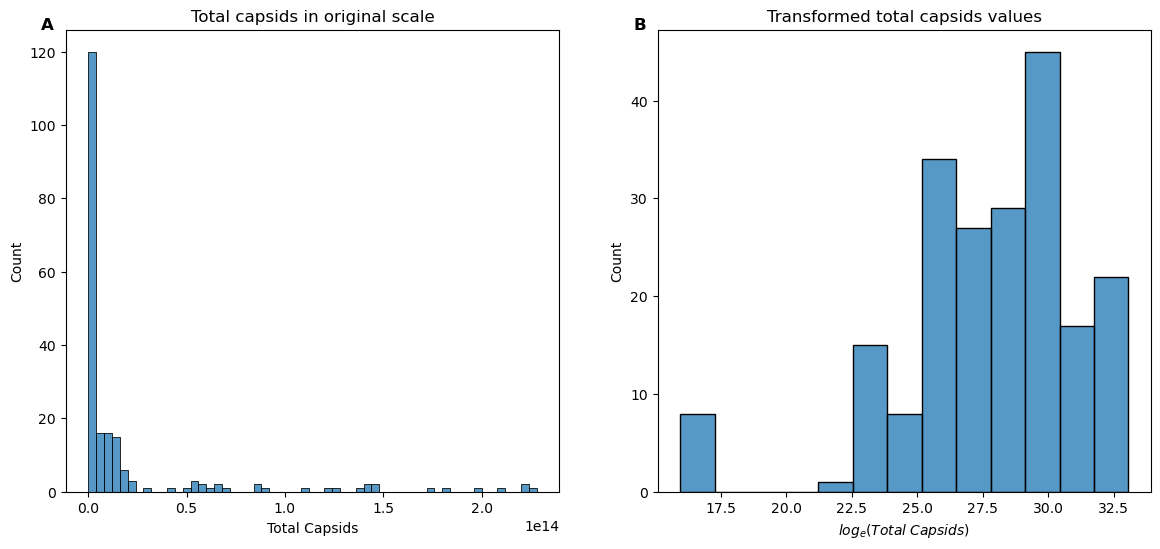

***Figure S1.*** *Total capsid titer in* ***(A)*** *normal and* ***(B)*** *log scale.*

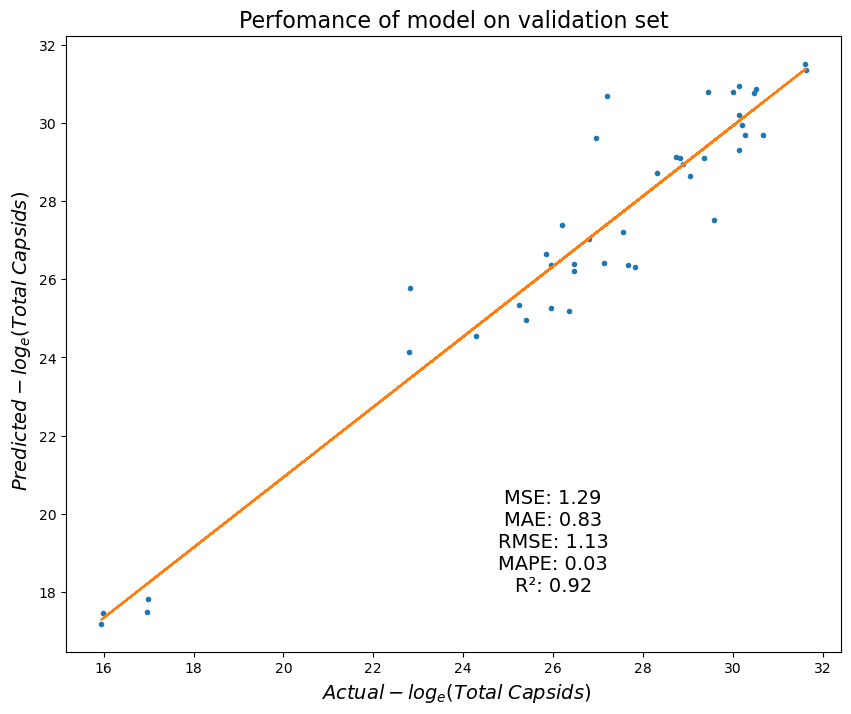

***Figure S2.*** *Comparison of the corresponding values predicted by the Gaussian process model with the Actual log(Total Capsids). The plot also includes annotations for the model's performance metrics on the hold-out validation set, which demonstrate low error and high accuracy. The acronyms MSE, MAE, RMSE, and MAPE stand for mean squared error, mean absolute error, and mean average percentage error, respectively.*

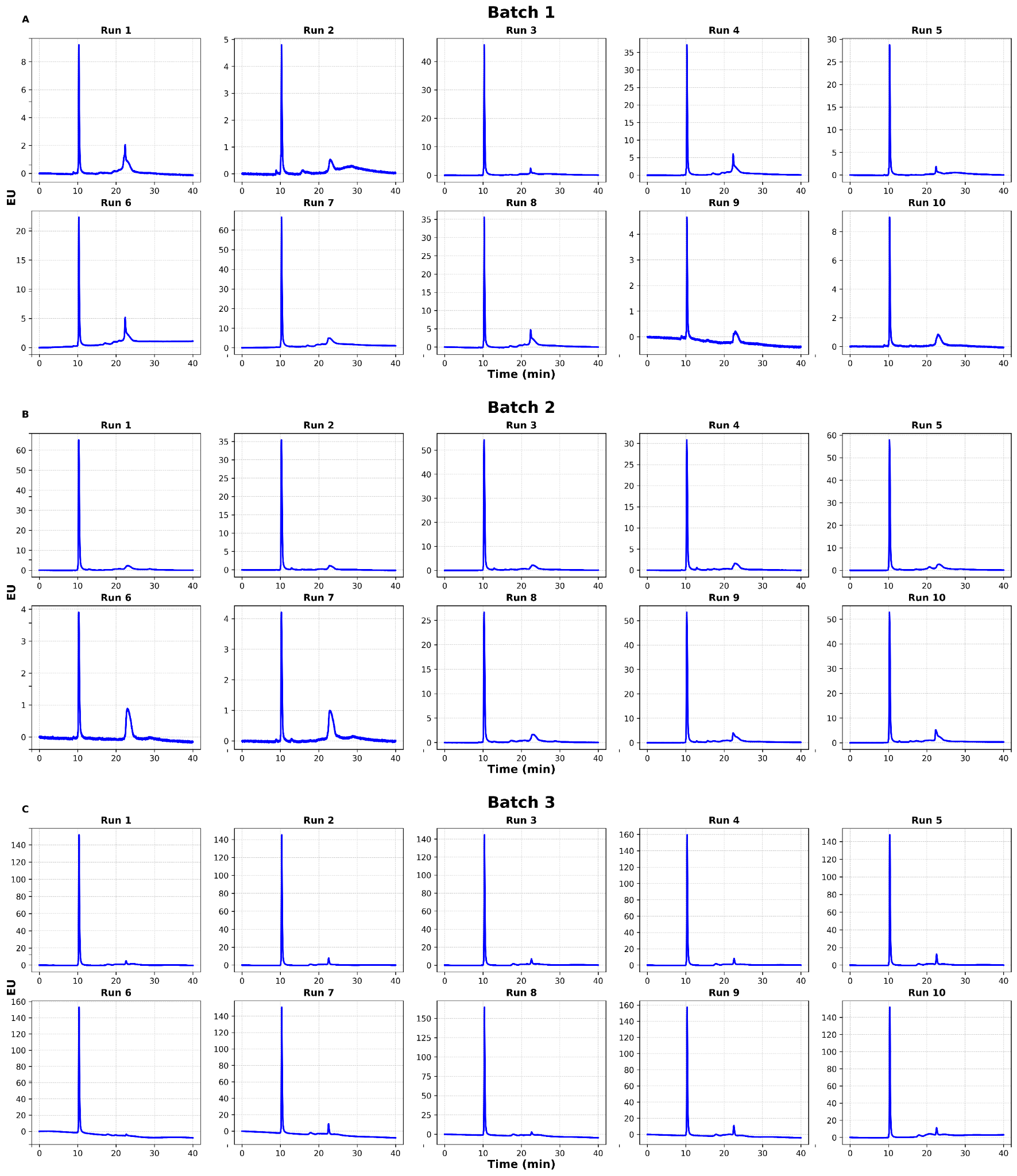

***Figure S3.*** *SEC-HPLC analysis of the elution fractions obtained by purifying AAV2 from HEK293 cell lysates using AAVidity operated with process input parameters selected across three iterations of the closed-loop Bayesian optimization process.*

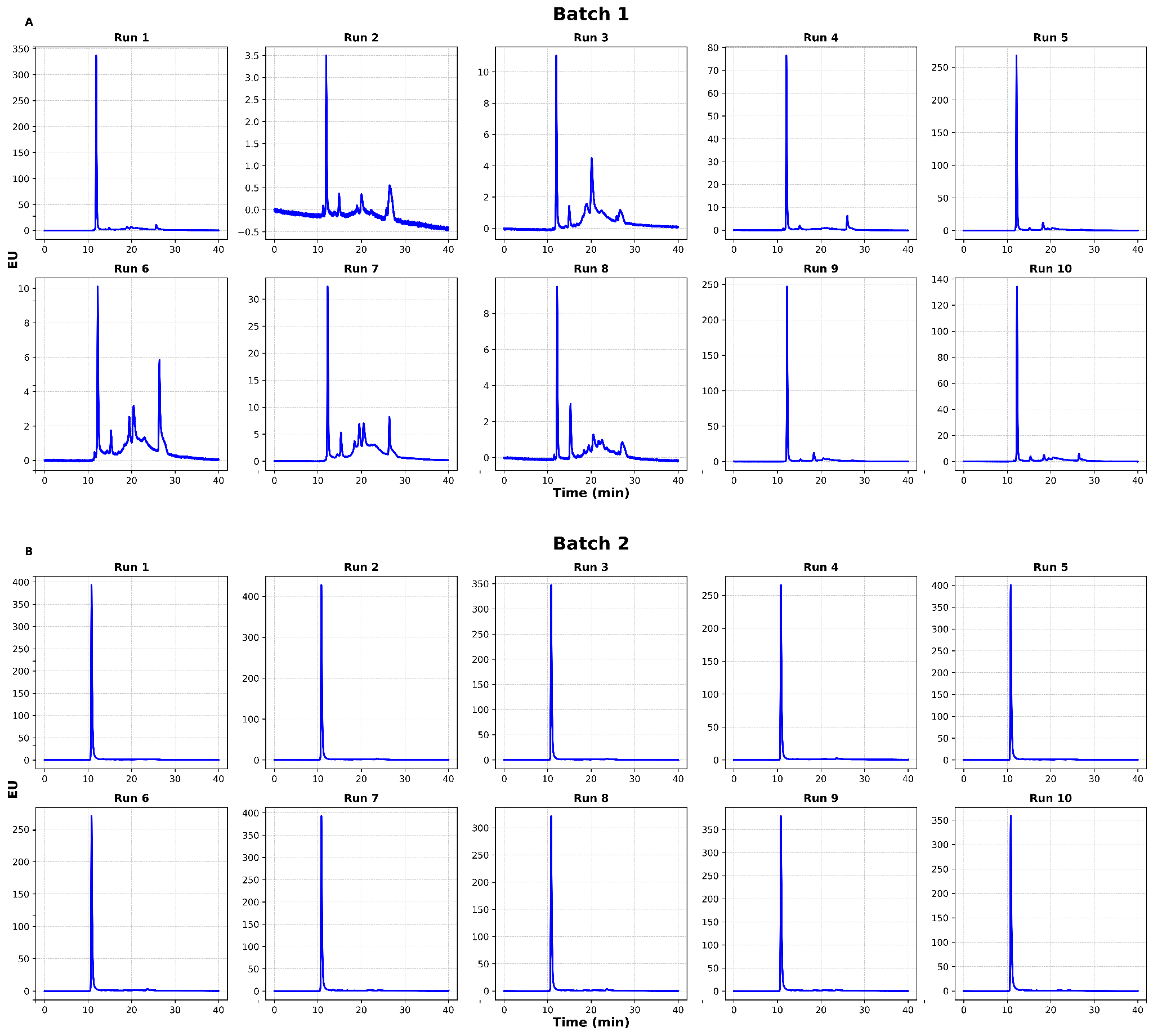

***Figure S4.*** *SEC-HPLC analysis of the elution fractions obtained by purifying AAV9 from HEK293 cell lysates using AAVidity operated with process input parameters selected across three iterations of the closed-loop Bayesian optimization process.*

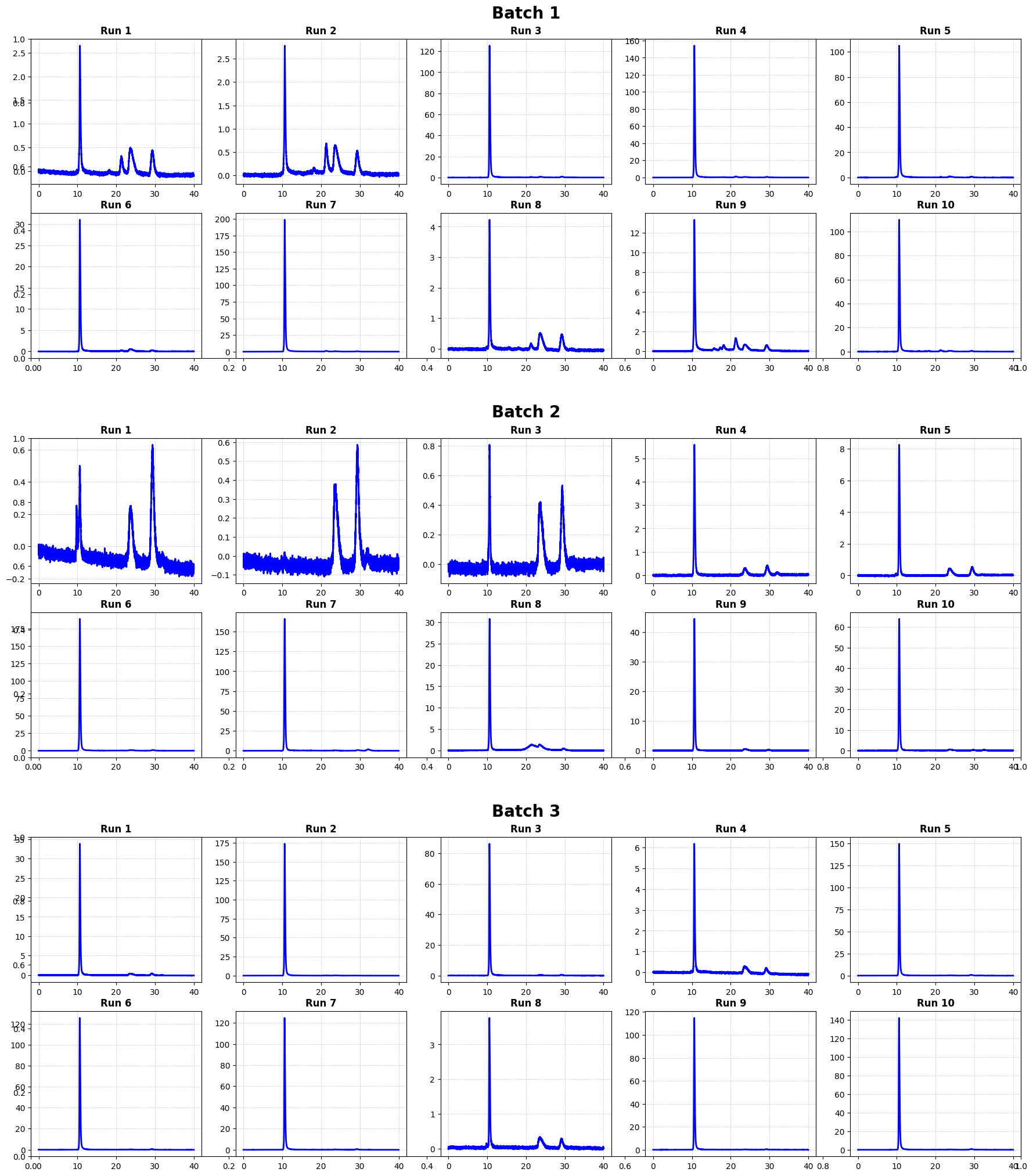

***Figure S5.*** *SEC-HPLC analysis of the elution fractions obtained by purifying AAV9 from HEK293 cell lysates using AAVidity operated with process input parameters selected across three iterations of the closed-loop Bayesian optimization process.*

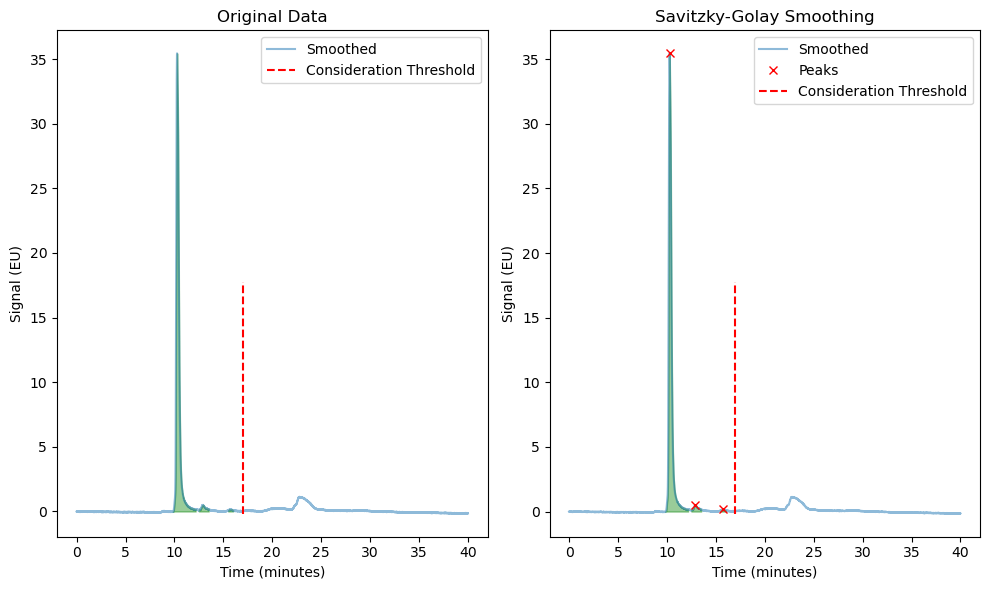

***Figure S6.*** *Integration of the peaks in the SEC-HPLC chromatograms either (A) before or (B) after Savitzky-Golay data smoothing for the calculation of the capsid purity.*
